# Discovery of photosynthesis genes through whole-genome sequencing of acetate-requiring mutants of *Chlamydomonas reinhardtii*

**DOI:** 10.1101/2021.02.17.431526

**Authors:** Setsuko Wakao, Patrick M. Shih, Katharine Guan, Wendy Schackwitz, Joshua Ye, Robert M. Shih, Mansi Chovatia, Aditi Sharma, Joel Martin, Chia-Lin Wei, Krishna K. Niyogi

**Author notes:** Jackson Lab, Farmington CT, 06032.

## Abstract

Large-scale mutant libraries have been indispensable for genetic studies, and the development of next-generation genome sequencing technologies has greatly advanced efforts to analyze mutants. In this work, we sequenced the genomes of 660 *Chlamydomonas reinhardtii* acetate-requiring mutants, part of a larger photosynthesis mutant collection previously generated by insertional mutagenesis with a linearized plasmid. We identified 554 insertion events from 509 mutants by mapping the plasmid insertion sites through paired-end sequences, in which one end aligned to the plasmid and the other to a chromosomal location. Nearly all (96%) of the events were associated with deletions, duplications, or more complex rearrangements of genomic DNA at the sites of plasmid insertion, and 1405 genes in total were affected. Functional annotations of these genes were enriched in those related to photosynthesis, signaling, and tetrapyrrole synthesis as would be expected from a library enriched for photosynthesis mutants. Systematic manual analysis of the disrupted genes for each mutant generated a list of 273 higher-confidence candidate photosynthesis genes, and we experimentally validated two genes that are essential for photoautotrophic growth, *CrLPA3* and *CrPSBP4*. The inventory of candidate genes includes 55 genes from a phylogenomically defined set of conserved genes in green algae and plants. Altogether, 68 candidate genes encode proteins with previously characterized functions in photosynthesis in *Chlamydomonas*, land plants, and/or cyanobacteria, 15 genes encode proteins previously shown to have functions unrelated to photosynthesis, and 190 genes encode proteins without any functional annotation, signifying that our results connect a function related to photosynthesis to these previously unknown proteins. This mutant library, with genome sequences that reveal the molecular extent of the chromosomal lesions and resulting higher-confidence candidate genes, represents a rich resource for gene discovery and protein functional analysis in photosynthesis.

## Introduction

Since the dawn of modern genetics, mutagenesis has been the primary vehicle to perturb the underlying genetic code of organisms, enabling scientists to investigate the genetic determinants underpinning biological systems. In the case of photosynthesis, much has been learned through mutagenesis of the unicellular green alga, *Chlamydomonas reinhardtii*, which has proven to be an indispensable reference organism for investigating the molecular components, regulation, and overall processes of photosynthesis (1, 2). *Chlamydomonas* has a haploid genome and an ability to use acetate as a sole carbon source, which facilitates the isolation and analysis of knock-out mutants that are defective in photosynthesis (3). Moreover, the advantage of working with a unicellular alga rather than a whole plant has facilitated the speed with which molecular and genetic studies can be carried out (4). Thus, the development of resources and tools to increase the breadth and depth of genetic studies in *Chlamydomonas* has advanced our ability to understand the molecular basis of photosynthesis.

Numerous large-scale mutagenesis and screening experiments have been carried out in *Chlamydomonas*, with some of the earliest efforts described over half a century ago (3,5,6). Classical mutagenesis studies have utilized chemical and physical mutagens, which induce untargeted genomic lesions and rearrangements across the genome. Identifying the causative mutations requires genetic mapping through crosses, an approach that is robust but time consuming. Insertional mutagenesis approaches, in which a selectable marker is transformed and randomly integrated into the genome, have facilitated molecular analysis, and many PCR-based techniques have been successfully employed in *Chlamydomomas* to rapidly identify flanking sequence tags (FSTs) from the site of marker insertion (7–14). However, the efficiency of FST recovery can be low (7) because of the complexity of events accompanying plasmid insertion such as concatemerization, chromosomal deletion or rearrangement, loss of the primer annealing sites, as well as difficulties with PCR from the *Chlamydomonas* nuclear genome, which is GC-rich and contains a high degree of repetitive sequences (15). High-throughput FST recovery has been achieved in *Chlamydomonas* (8, 10) and has offered a large collection of insertional mutants for the scientific community while enabling large-scale mutant analysis of photoautotrophic growth (9).

The advent of next-generation sequencing methods has dramatically improved our ability to identify mutations by whole-genome sequencing (WGS). In *Chlamydomonas*, this approach was initially combined with linkage mapping to identify point mutations in flagellar mutants (11, 12), and it was used subsequently for point mutations affecting the cell cycle (13, 14) and light signaling (16, 17). In the case of insertional mutants, WGS has been used extensively to identify insertion sites in bacteria and some microbial eukaryotes with smaller genomes (18–20) but only for a relatively small number of mutants in *Chlamydomonas* (21). In maize, due to its large genome, high-throughput next-generation sequencing of *Mu* transposon insertion sites has been applied only after enrichment for the transposon sequence (22), whereas the large volume of insertion site information of T-DNA insertion lines in *Arabidopsis* was obtained from traditional PCR-based FST isolation (23–25).

We have previously generated a large insertional mutant population of *Chlamydomonas* by transformation with a linearized plasmid conferring paromomycin or zeocin resistance, and we identified mutants with photosynthetic defects (*i.e.*, acetate-requiring and/or light-sensitive and reactive oxygen species-sensitive mutants) (7, 26). However, we were only able to obtain FSTs for 17% of the mutants using PCR-based approaches. Here we employed low-coverage WGS of a subset of 660 mutants to identify the plasmid insertion sites and accompanying structural variants, and we found 1405 genes that are affected by the plasmid insertion in 509 mutants. We generated a list of 273 genes from 348 mutants that we refer to as higher-confidence causative genes, enabling the discovery of 205 potential photosynthesis genes; 190 genes of previously unknown function and 15 genes previously shown to have functions unrelated to photosynthesis. We experimentally validated two genes, *CrLPA3* and *CrPSBP4,* that are required for photoautotrophic growth in *Chlamydomonas*. In addition, our data provide insight into the spectrum of mutations that are induced by insertional mutagenesis in *Chlamydomonas*.

## Results

### Identification of insertion sites by mapping of discordant read pairs

We re-screened our *Chlamydomonas* photosynthetic mutant collection (7, 26) for growth on minimal and acetate-containing media under three light conditions (dark, D; low light of 60-80 µmol photons m^-2^ s^-1^, LL; and high light of 350-400 µmol photons m^-2^ s^-1^, HL) and for maximum photochemical efficiency of photosystem (PS) II (F_v_/F_m_) (S1 Table). An example of the phenotyping is shown in Figure 1. A total of 660 mutants, most of them with a growth phenotype and with resistance to either zeocin or paromomycin, indicative of the presence of the linearized plasmid sequence used for insertional mutagenesis, were chosen for WGS and herein will be referred to as the Acetate-Requiring Collection (ARC).

**Fig 1.**
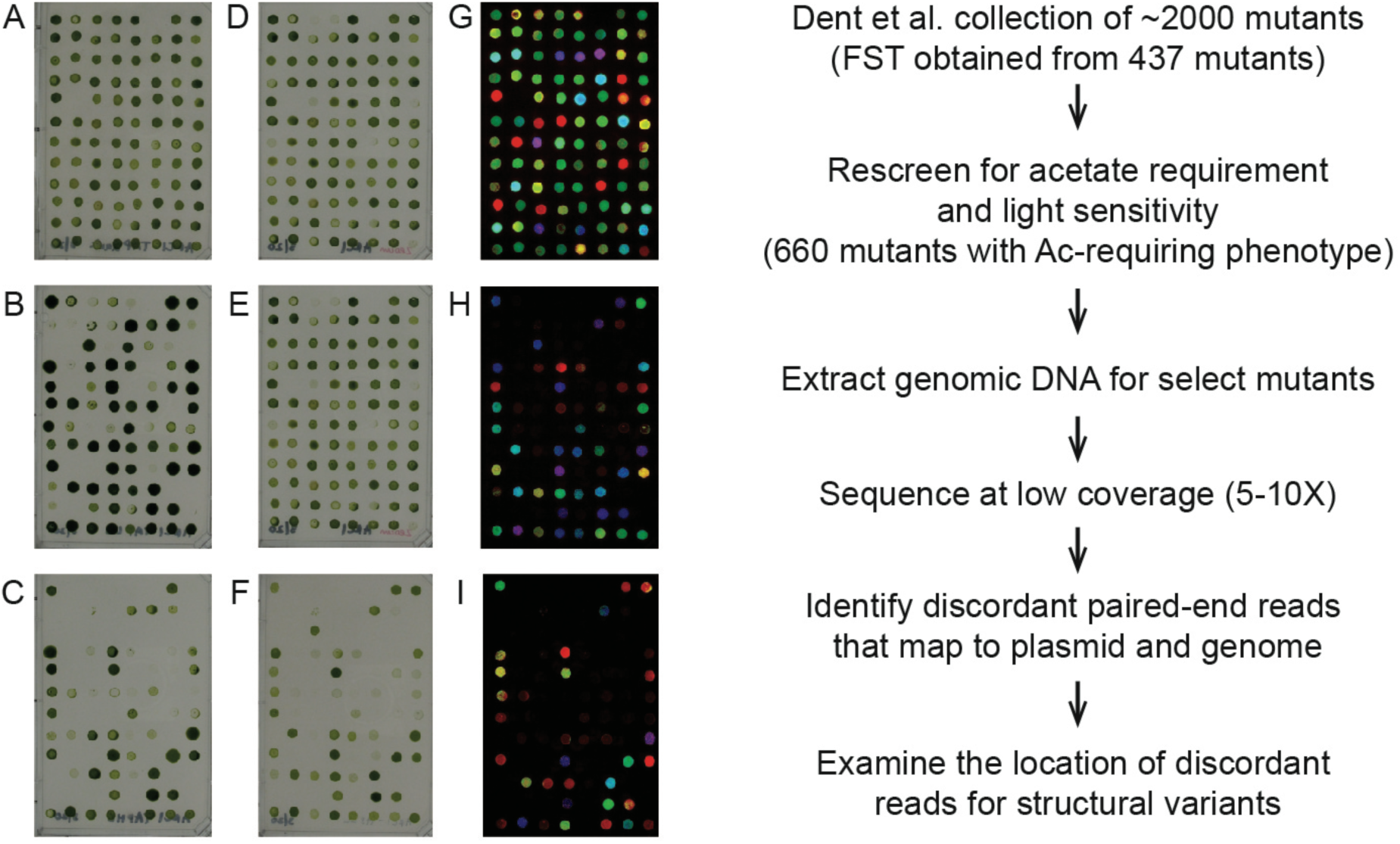
Growth and chlorophyll fluorescence screen pipeline. Mutants were scored for growth on (A) D+ac, (B) LL+ac, (C) HL+ac, (D) LL+ac+zeocin, (E) LL-min, (F) HL-min. F_v_/F_m_ values were measured on cells grown on (G) D+ac, (H) LL-min, (I) HL-min. FST, flanking sequence tag. A representative plate spotted from a 96-well plate is shown. D, dark; LL, low light; HL, high light; +ac, added acetate; min, minimal media.

Genomic DNA was extracted from the 660 ARC mutants and submitted for low-coverage, paired-end WGS with a target depth of sequence coverage for each mutant between 5 and 10. The average sequencing depth across samples was 7.44. Paired-end reads that showed one end mapping to the plasmid used for mutagenesis and the other to a chromosome location were used to identify the plasmid insertion site(s) in each mutant. Plasmid insertion sites were not identified for 72 mutants, because few plasmid sequence reads were detected or the other end mapped to a low complexity region of the *Chlamydomonas* genome. 79 mutants had insertions that were not unique within the population (33 were duplicated, three were triplicated and one was quadruplicated) and were removed from further analysis. The remaining 509 mutant sequences were further analyzed for structural variants (insertions, deletions, and rearrangements) that occurred during insertional mutagenesis.

Figure 2 illustrates the types of structural variants detected by analysis of the paired-end sequence data. Most sequence read pairs were concordant, i.e., they showed the expected orientation and distance with respect to each other when mapped to the *Chlamydomonas* genome (Figure 2, dark gray arrows). In contrast, discordant pairs showed the incorrect orientation or distances that were closer or further from each other than expected based on the genome fragmentation that was performed during sequencing library preparation (genomic DNA was sheared to approximately 600 bp) or on different chromosomes. In Figure 2, the discordant reads are shown as colored arrows, with each color representing a chromosome (or plasmid) to which the corresponding paired-end read was mapped. Each of these genomic sites where sequence read pairs were discordant is listed in S1 Table as a “Discordant site”.

**Fig 2.**
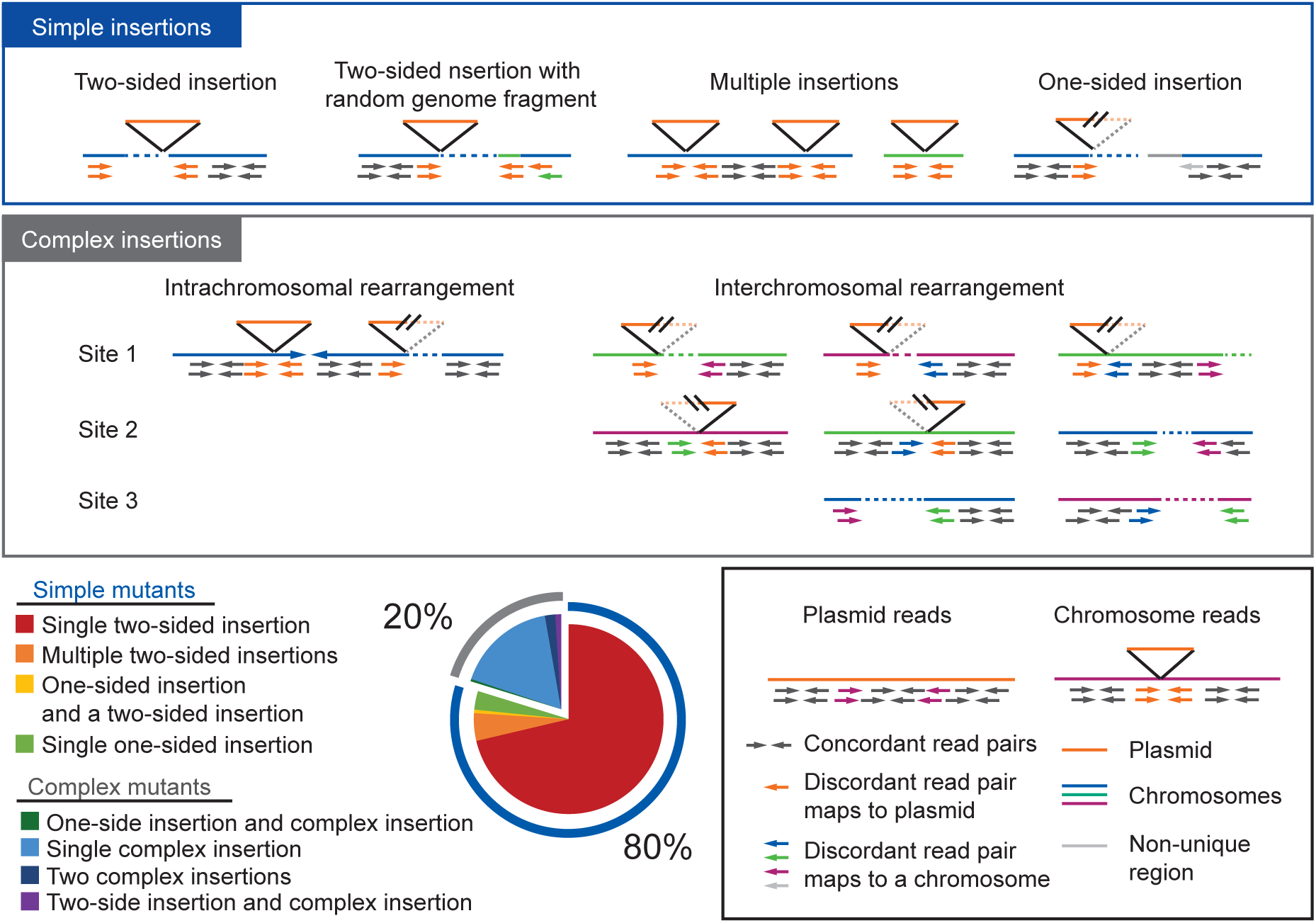
Examples of structural variations and the frequency mutants with simple or complex insertions in ARC. Boxes contain schematic examples of mapped reads as seen in IGV. Black box, mapped reads (concordant and discordant) against plasmid and chromosome. Blue box, examples of “Simple insertions”; Gray box, examples of “Complex insertions”. Gray box shows examples of different complex insertions that are intra- or interchromosomal rearrangements. Second from left in gray box shows a possible translocation between two chromosomes. Pie chart shows frequency of “Simple mutants” containing only simple insertions and “Complex mutants” containing complex insertions.

At most of the plasmid insertion sites, two sets of discordant read pairs were found, with their chromosomal reads oriented toward each other and their paired-end reads mapping to the plasmid sequence (Figure 2 blue box). We refer to these 425 events as two-sided insertions, where both sides of the plasmid insertion were unambiguously mapped (S1 Table, column “Number of sides paired with plasmid at site”, 2). Another large group of discordant sites displayed only one set of discordant read pairs located on one side of the plasmid insertion (referred to as one-sided insertions Figure 2; S1 Table, column “Number of sides paired with plasmid at site”, 1). The read-pairs on the other side of the plasmid insertion could not be mapped in 21 of these insertion sites because (i) it was at a repetitive region (14 mutants) and (ii) it had no discordant reads (7 mutants). These 21 one-sided insertions together with the 425 two-sided insertions making a total of 446 insertions and were considered to be simple insertions (S1 Table). In the rest of the one-sided insertions, the other side of the plasmid insertion paired with another chromosomal region indicating an occurrence of a more complex chromosomal rearrangement. Insertions that paired with another chromosomal location was considered a complex insertion. The frequencies of two-sided, one-sided, and complex insertions are shown in S1 Figure.

A total of 406 mutants (80%) contained only simple insertions accounting for 435 out of the total 446 simple insertions (11 mutants contained both simple and complex insertions) (Figure 2, Simple mutants). Among these 406 mutants, 24 mutants had multiple (two or three) two-sided insertions accounting for 50 insertions, and three mutants had one two-sided insertion and one one-sided insertion (Figure 2, S1 Figure). In 17 mutants, the multiple simple insertions occurred on the same chromosome, and six of these had tandem two-sided insertions that disrupted the same or neighboring genes. In 10 two-sided insertions (∼1.8%), there appeared to be a short random fragment of another chromosome inserted together with the plasmid (Figure 2, Two-sided insertion with random genome fragment). The original locus of these random fragments did not show a lack of mapped sequence reads but rather showed double the abundance of reads mapping to the small region, indicating that it was an extra copy of the same sequence at the insertion site, similar to what was observed in a previous study but at a lower frequency in ARC (8).

The other group of 103 mutants (20%) contained at least one complex insertion (Figure 2 “Complex mutants”; also see S1 Table, “Pairing with other discordant site(s) of the same mutant”). Nine of these mutants had a coexisting two-sided insertion, two mutants had an additional one-sided insertion, and five mutants contained two independent complex insertions. Some of these rearrangements occurred on a single chromosome, and others involved two or more chromosomes (Figure 2, gray box). Among interchromosomal rearrangements, 13 of them involved two one-sided insertions that were paired to each other (Figure 2 gray box). These together may represent chromosomal translocation events resulting in two chimera chromosomes. In all of these possible translocation events, the plasmid sequence was present in one junction and not in the other. The proportion of complex insertion events was similar among the three plasmids used for transformation (pSP124S, pMS188, and pBC1). Validation of these complex structural variants would require *de novo* assembly of sequencing reads. Most mutants only contained only two-sided or only complex insertions; 387 mutants (76%) had only two-sided insertion(s) (Figure 2, red and orange slices), 92 had only complex insertion(s) (18%) (Figure 2, light blue slice), and only a small proportion of mutants contained a mix of two-sided, one-sided, or complex insertions.

**S1 Fig.**
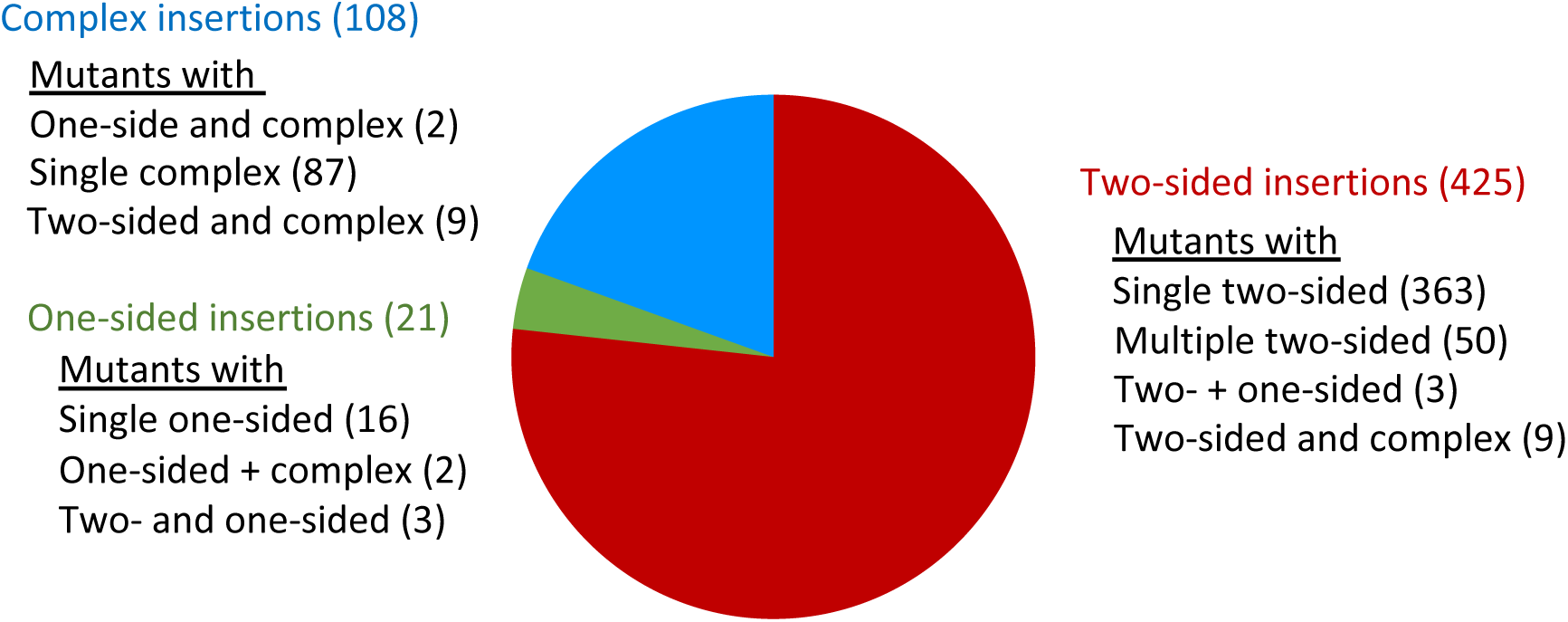
Proportion of different types of insertions observed in ARC. The frequency of the different types of insertions. Some insertions coexist with another insertion in a mutant. The number of mutants grouped by the types of insertions it contains is listed along with the number of insertions accounted for in that group.

In summary, low-coverage WGS data for 509 ARC mutants identified 406 mutants that contained only simple insertions accounting for 435 out of 446 total simple insertions, whereas 103 mutants contained complex insertions that were associated with chromosomal rearrangements such as inversions and translocations.

### Analysis of deletions and duplications associated with insertional mutagenesis

Insertional mutagenesis in *Chlamydomonas* has been previously associated with deletions and duplications at the site of plasmid insertion, especially when using glass bead for transformation (e.g. *cpld38*, *cpld49*, *npq4, rbd1*) (27–29). Focusing on the 425 two-sided insertions, we found deletions associated with 374 insertions (88%). A wide range of deletion sizes was observed, with a bimodal distribution peaking at 101-1000 bp and 10 - 100 kb when plotted at log_10_-scale, the largest deletion being 133 kb (Figure 3A). Duplications occurred less frequently (7%), in a total of 29 insertion events (Figure 3B), and all were less than 1000 bp. Perfect insertions lacking any duplications or deletions were found in only 22 events (5%). Despite the high frequency and relatively large size of many deletions, more than half (220 insertions) of the entire set of 425 two-sided insertions affected only a single gene (Figure 3C).

**Fig 3.**
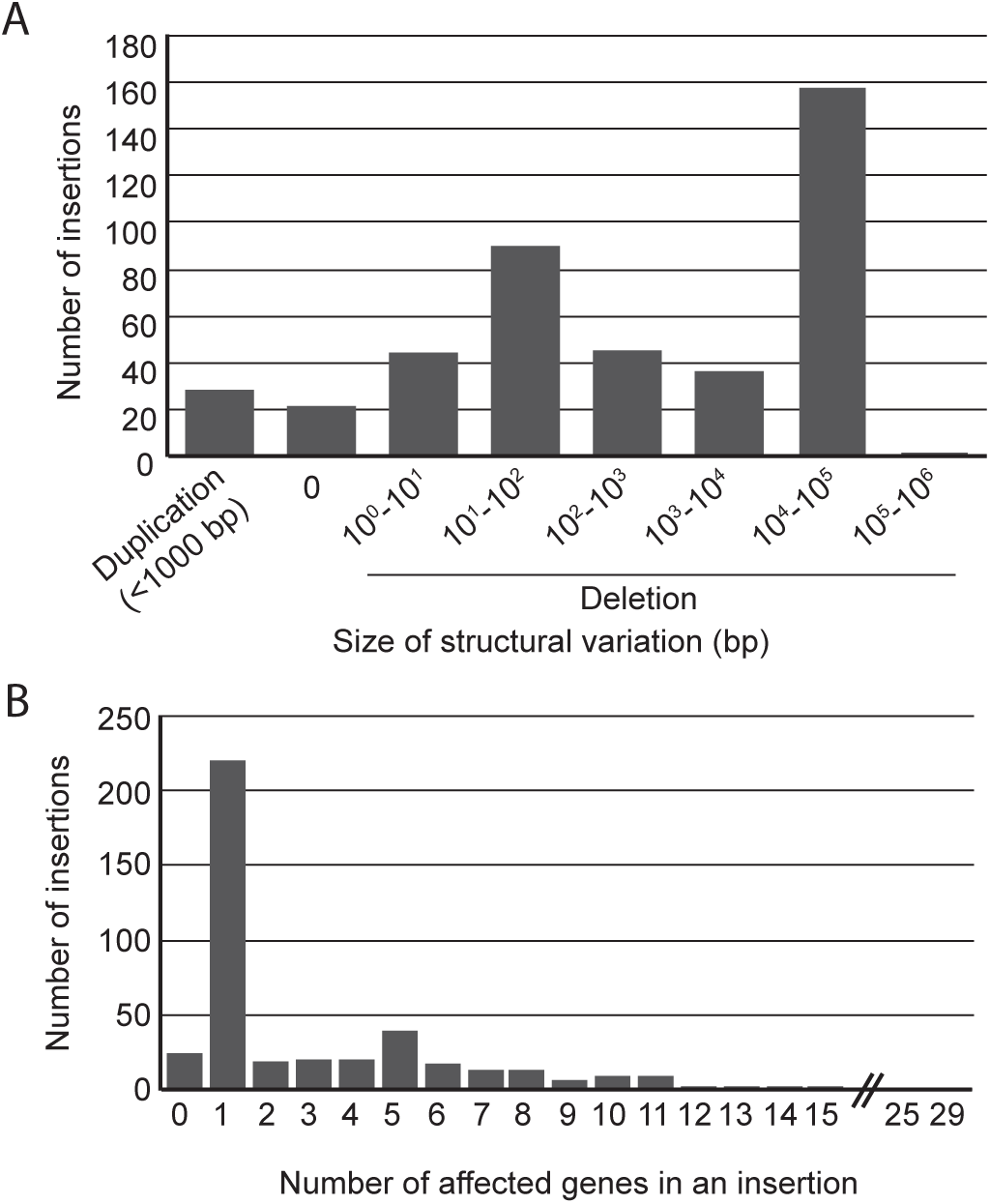
Structural variation accompanying insertions. (A) Duplication and deletion sizes and (B) number of mutants grouped by the number of genes affected by two-sided insertions. Only two-sided insertions were included in this analysis.

### Genetic linkage between acetate-requiring phenotype and antibiotic resistance

To determine if the phenotype of ARC mutants was likely caused by the plasmid insertion, we back-crossed 89 mutants to the wild type (WT) and analyzed the genetic linkage of the acetate-requiring phenotype and antibiotic (paromomycin) resistance in the respective progenies. The acetate-requiring phenotype was closely linked to the antibiotic resistance in 88% (77 out of 88 that produced viable zygospores) of mutants that were tested (S1 Table, column “Genetic Linkage”). In each cross, approximately 100 zygospores were collected and tested for recombination between the acetate-requiring phenotype and paromomycin resistance by selecting for progeny that were able to grow on minimal medium with paromomycin (S2 Figure). The lack of recombination and therefore growth indicates that the genetic distance between the mutation causing the acetate-requiring phenotype and paromomycin resistance is less than 0.5 cM, estimated to be 50 kb on average in the *Chlamydomonas* genome (15).

### Identification of secondary mutations using WGS data

In addition to the deletions associated with plasmid insertions in the ARC mutants, we searched for and found 68 other deletions using Pindel (30) (S2 Table). The size of the deletions ranged from 20 bp to 36 kb, with a majority of them (55 deletions) being less than 100 bp (S2 Table). The deletions were visually confirmed on alignments as direct gaps in reads and/or the lack of reads within the region, depending on the size. This was not expected to be an exhaustive search for such deletions. For example, low-coverage regions could be difficult to distinguish from a deletion. Nevertheless, some of the deletions affected clear candidate genes that could be responsible for the mutant phenotype. For example, the CAL014_01_19 mutant was found to contain a 21-bp deletion in Cre01.g013801, a GreenCut2 gene (conserved within genomes of land plants and green algae but absent from non-photosynthetic organisms (15, 31)) annotated as a tocopherol cyclase (*VTE1*). The deletion occurred at the junction of intron 7 and exon 8, which could affect splicing and translation of a functional protein (S3 Figure). Because tocopherols are important for photoprotection in *Chlamydomonas* (32) disruption in the *VTE1* gene could explain this mutant’s high light-sensitive phenotype (S1 Table). In support of this hypothesis, a second mutant in the ARC, CAL033_02_19, had a 33-bp deletion in this locus. Interestingly, this mutant has a less severe phenotype (S1 Table), consistent with the plasmid insertion and deletion positioned in the 3’-UTR of the gene, which may have led to a partial loss of function (S3 Figure).

**S2 Fig.**
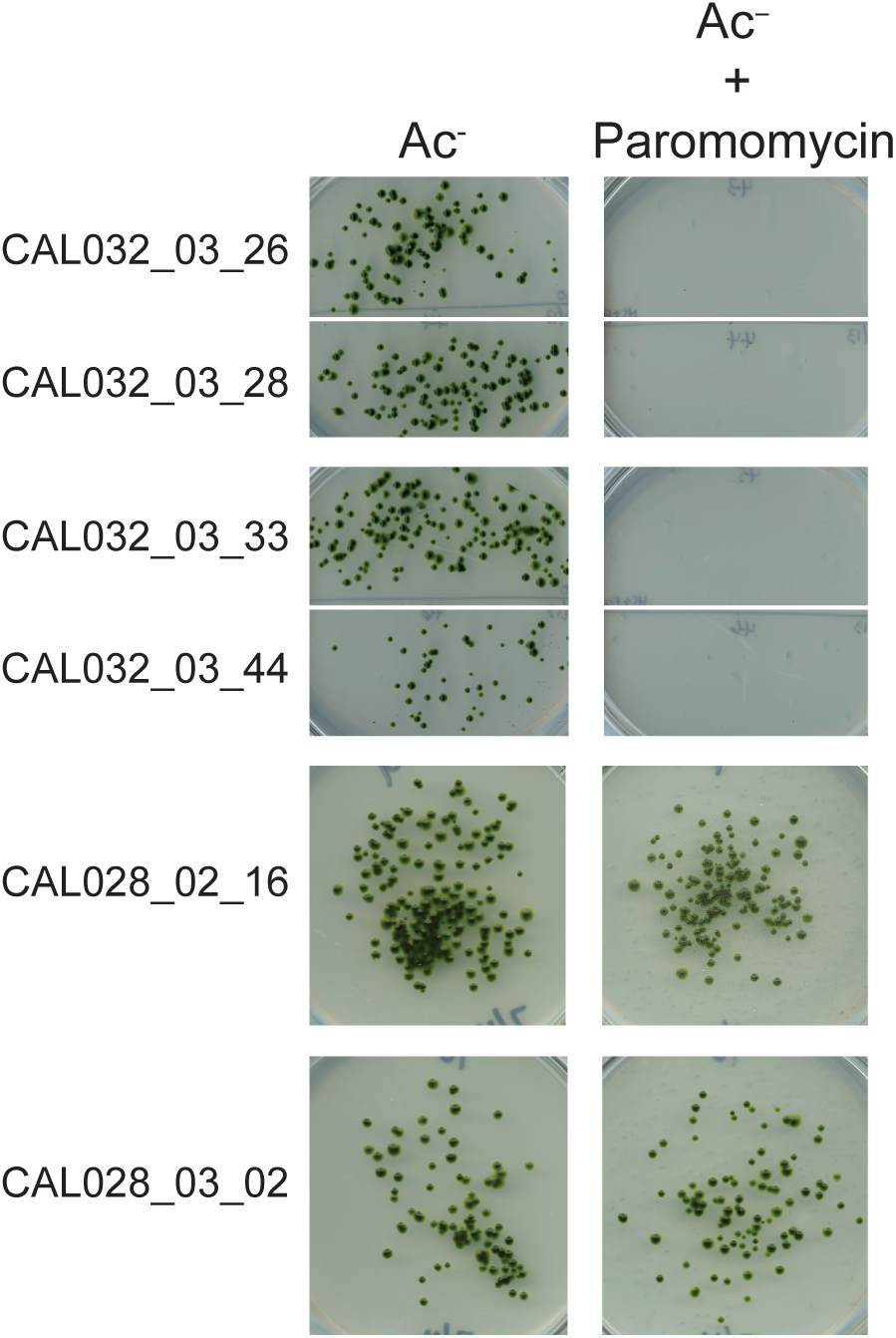
Genetic linkage test of par^R^ and Ac^−^ phenotypes. Mutants (ac^−^ par^R^) were crossed with WT (AC^+^ par^S^) cells and the zygospores were tested for growth on minimal media with and without paromomycin. Absence of growth on min+paromomycin indicates the genetic linkage of the two phenotypes.

Among the 11 mutants whose acetate-requiring phenotype did not cosegregate with its paromomycin resistance, one (CAL036_02_12) had a strong acetate-requiring phenotype (S1 Table) and contained a 36-kb deletion located 2 Mb away from the plasmid insertion on chromosome 7. This resulted in a deletion of seven genes (Cre07.g346050, Cre07.g346100, Cre07.g346150, Cre07.g346200, Cre07.g346250, Cre07.g346300, and Cre07.g346317). One of these (Cre07.g346050) is *COPPER RESPONSE DEFECT 1* (*CRD1*), and *crd1* mutants have a conditional phenotype, lacking accumulation of PSI only under copper deficiency (33). Another mutant (CAL029_03_36) has a one-sided insertion in *CRD1* and was only modestly affected in growth in HL (S1 Table), suggesting that the loss of CRD1 is not the cause of the severe phenotype of CAL036_02_12. Another one of the deleted genes is annotated as phytol kinase (Cre07.g346300). Chlorophyll degradation and phytol remobilization through phytol kinase (*VTE5*) and phytol phosphate kinase (*VTE6*) are important for *α*-tocopherol biosynthesis and their disruption results in high light sensitivity in tomato (34) and *Arabidopsis* (35). The light sensitivity observed in CAL036_02_12 is similar to that of tomato plants silenced for *VTE5* (34) and strongly suggests that Cre07.g346300 is the causative gene for the mutant phenotype. The remaining 10 mutants whose acetate-requiring phenotype is unlinked to the plasmid insertion would be candidates for higher-coverage WGS to search for causative mutations.

**S3 Fig.**
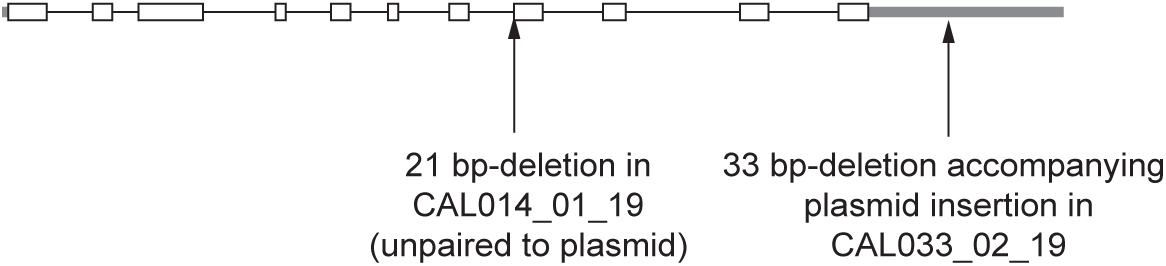
Two mutant alleles in tocopherol cyclase (Cre01.g013801) in ARC. Schematic representation of the disruption sites in CAL014_01_19 a strictly acetate-requiring mutant and CAL032_02_19, a mutant with comparatively moderate phenotype.

### Genes with multiple mutant alleles in the ARC

In total, 1405 genes were directly affected by the 554 plasmid insertions in 509 mutants. There are many more affected genes compared to the number of mutants from which they originate due to disruption of multiple genes by large deletions. S3 Table lists all of the disrupted genes and their available annotations.

To begin identifying causative mutations, we searched for genes that were affected in multiple ARC mutants. Figure 4A shows the number of alleles of the 1405 genes that occur in the ARC. Interruption/deletion of 1053 genes only occurred once, while 212 genes have two alleles and 94 genes have three alleles. Some genes appeared on the list of affected genes more than three times (Figure 4A). However, because disruption of multiple genes occurred in approximately half of the ARC mutants, many of these genes represented by multiple alleles are likely not causative for the mutant phenotype. Some of the genes appear more frequently on the list simply because of their proximity to the causative gene. Figure 4B shows an example of such an occurrence for *CPSFL1* (Cre10.g448051). Seven ARC mutants had deletions ranging from 22 to 130 kb in a region on chromosome 10 (CAL028_01_03, CAL033_04_04, CAL031_01_04, CAL039_03_10, CAL007_02_07, CAL038_02_20, and CAL028_01_06) (S1 Table). 33 genes were affected by the deletions in these mutants, including seven genes affected in all seven mutants, which makes it difficult to narrow down to a single causative gene. One additional mutant (CAL29_02_48) had a complex insertion event involving four different chromosomes, but strikingly it shared a single affected gene (*CPSFL1*, containing a 10-bp deletion) with the other seven mutants. All eight mutants exhibited a strict acetate-requirement and severe light-sensitivity phenotype (S1 Table), and in-depth characterization of the CAL028_01_06 mutant showed that *CPSFL1* is involved in carotenoid accumulation and is essential for photoautotrophic growth in *Chlamydomonas* and *Arabidopsis* (36, 37).

**Fig 4.**
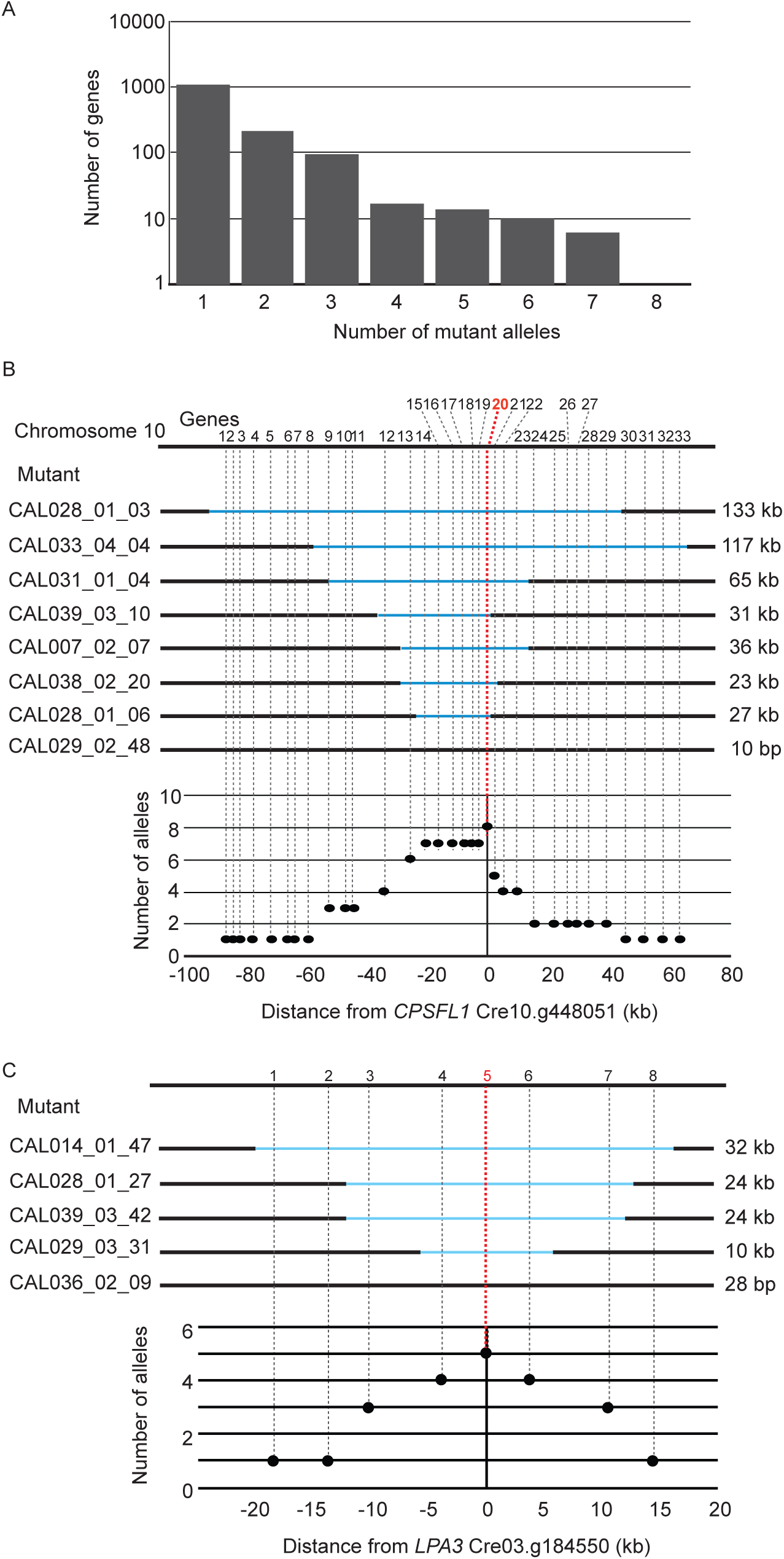
Genes represented by multiple mutant alleles are more likely to be causative genes. (A) Number of genes among all 1405 genes affected in ARC grouped by the number of mutant alleles that represent the gene. Schematic of mutant alleles disrupted in (B) *cpsfl1* mutants and (C) *lpa3* mutants and the allele frequencies of surrounding genes. Note that not all genes with multiple mutant alleles are causative; some occur among the 1405 affected genes because of their physical proximity to the true causative genes.

The *CrLPA3* gene (Cre03.g184550, hereon *LPA3*) is another example of a gene that was affected in multiple mutants (Figure 4C). The CAL014_01_47, CAL028_01_27, CAL039_03_42, CAL029_03_31, and CAL036_02_09 mutants had overlapping deletions ranging from 28 bp to 32 kb in the same region on chromosome 3, and all five mutants exhibited a strict acetate-requiring phenotype in HL (S1 Table). By comparing the disruption frequencies, we identified *LPA3* as the only gene that was affected in all five mutants.

### *LPA3* and *PSBP4* are essential for photoautotrophic growth

We proceeded to validate the WGS data and identify two genes as necessary for photoautotrophic growth in *Chlamydomonas*. In one case (*LPA3*), multiple alleles were present in the ARC, whereas only a single allele of the other gene *CrPSBP4* (hereon *PSBP4*) was present. Three *lpa3* mutants (CAL028_01_27, CAL039_03_42, and CAL040_01_25) were selected for further analysis (and renamed as *lpa3-1*, *lpa3-2*, and *lpa3-3*, respectively) The WGS data indicated that the *lpa3-1* and *lpa3-2* mutants had very similar deletions of 24 kb that affected the same five genes (S1 Table). The deletion was confirmed by amplifying genomic regions across the predicted deletion by PCR in both mutants (Figure 5A), although it was not possible to amplify the plasmid sequence at the site of the deletion. The *lpa3-3* mutant was predicted from WGS to have a 4-bp deletion and plasmid insertion in the 5’-UTR of Cre03.g184550, which was confirmed by sequencing a PCR fragment of the region from the mutant (Figure 5A), but it was not included in S1 Table, because it was one of the 79 mutants with a non-unique insertion site (see above in section “Identification of insertion sites by mapping of discordant read pairs”). All three mutants had an acetate-requiring phenotype (Figure 4B). The gene Cre03.g184550 encodes a GreenCut2 protein (CPLD28) (31), and is annotated as an ortholog of *Arabidopsis* LOW PSII ACCUMULATION 3 (LPA3). *Arabidopsis* LPA3 has been reported to be involved in the assembly of photosystem II (38), although the publication on the function of this protein was later retracted (39). Complementation with a genomic DNA clone of Cre03.g184550 (*LPA3*) including 1.2 kb upstream of the transcription start site rescued all three mutants, demonstrating that the disruption of this gene was responsible for the acetate-requiring phenotype of these mutants. Mutants lacking LPA3 exhibited very low F_v_/F_m_ values even in the dark (Figure 5C). This suggests that *Chlamydomonas* LPA3 is required for the assembly of PSII even in the absence of light, resulting in a much more severe phenotype than *lpa3* single mutants in *Arabidopsis*, which were able to grow in LL on soil (38). The low F_v_/F_m_ phenotype of the mutants was rescued in the complemented lines in all light conditions (Figure 5B, C).

**Fig 5.**
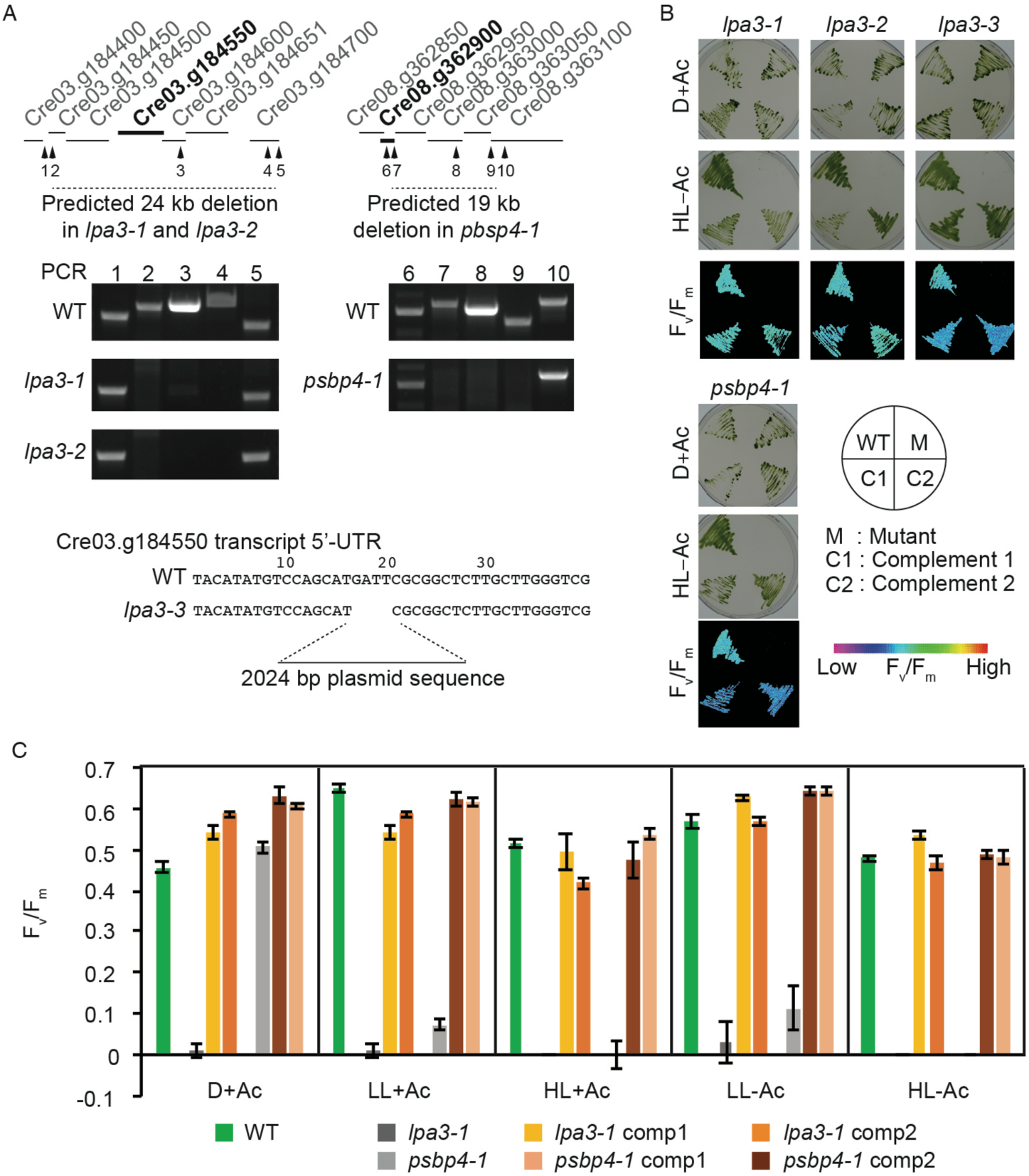
Identification of *CrLPA3* and *CrPSBP4* required for photoautotrophic growth. (A) Schematic of loci and deletions indicated from whole-genome sequence data in mutants *lpa3-1* (CAL028_01_27), *lpa3-2* (CAL039_03_42), and *lpa3-3* (CAL040_01_25) that share a disruption in Cre03.g184550, gene encoding a predicted ortholog of Arabidopsis LOW PHOTOSYSTEM II ACCUMULATION 3 (LPA3) and mutant *psbp4-1* (CAL032_04_48) that had a deletion encompassing Cre08.g362900, a gene encoding a protein predicted as PSBP4. Numbered arrowheads indicate the PCR probes used in testing for deletions shown in the agarose gel photos. WT and *lpa3-3* sequences indicate the plasmid insertion site and associated 4 bp-deletion. (B) Growth and chlorophyll fluorescence phenotype of WT, mutants and their complemented lines. Cells were grown with acetate in the dark, without acetate under 400 μmol photons s^-1^ m^-2^ and imaged for growth and F_v_/F_m_ measurements (HL-Ac). F_v_/F_m_ value are represented by false colors as shown in the reference bar. (C) F_v_/F_m_ values of each genotype under different growth conditions. comp, complemented line.

The mutant CAL032_04_48 (renamed as *psbp4-1*) required acetate for growth and exhibited light sensitivity even in the presence of acetate, and its F_v_/F_m_ was reduced compared to that of the WT when grown in the light (S1 Table, Figure 5C). Its WGS indicated two tandem simple insertions disrupting five genes. Among them, Cre08.g362900, annotated as encoding a thylakoid luminal PsbP-like protein (PSBP4), presented itself as a clear candidate to be the gene responsible for the phenotypes. The PSBP4 ortholog of Arabidopsis has been shown to involved in the assembly of PSI (40, 41). The deletion in *psbp4-1* was confirmed by PCR (Figure 5A), and the mutant phenotype was rescued by transforming with genomic DNA including Cre08.g362900 and upstream region, demonstrating that disruption of *PSBP4* was the cause of the acetate-requiring and light-sensitive phenotypes of this mutant (Figure 5B, C).

### Curation of higher-confidence photosynthesis candidate genes

To identify candidate genes that are likely to be responsible for the ARC mutant phenotypes, we focused on the 406 mutants with only simple insertions (Figure 2). We reasoned that a mutant with a simple insertion event is more likely to have a causative gene within its disrupted gene list than a mutant with a complex insertion event that is accompanied by large-scale chromosomal rearrangements, which could cause unpredictable changes in expression of neighboring genes due to alterations in promoters, enhancers, and chromatin environment. For each of the 406 mutants with simple insertions, we applied a series of criteria to generate a list of genes that are the strongest confidence candidates for being genes that are responsible for the ARC mutant phenotype. If a mutant contained a single, simple insertion that disrupts a single gene then that gene was immediately considered to be a higher-confidence candidate. If a mutant contained a simple insertion with multiple genes disrupted by an associated deletion, then we manually analyzed the genes and selected the best candidate, considering whether it was a GreenCut2 gene and/or whether it encoded a protein with annotation or domains indicating a possible function in photosynthesis (e.g. redox, chlorophyll *a/b*-binding, Fe-S cluster). 78 GreenCut2 genes that were disrupted in 509 ARC mutants (Table 1) and were considered strong candidates unless there was an even stronger candidate based on functional annotation. As was shown for *cpsfl1* (Figure 4B) and *lpa3* (Figure 4C), mutants with overlapping disrupted genes were also compared to find the strongest candidate (gene with highest disruption frequency). Neighboring genes that were co-disrupted with the strongest candidates were deemed non-candidates in all the mutants. As a final criterion, we searched candidate genes derived from analysis of other existing photosynthesis mutant libraries and identified overlaps with *Chlamydomonas* genes whose disruption affected photoautotrophic growth (9), orthologous genes from the maize Photosynthetic Mutant Library (PML, http://pml.uoregon.edu/pml_table.php) (42), and orthologous genes identified from Dynamic Environmental Photosynthetic Imaging (DEPI) of *Arabidopsis* mutants (43).

**Table 1.**
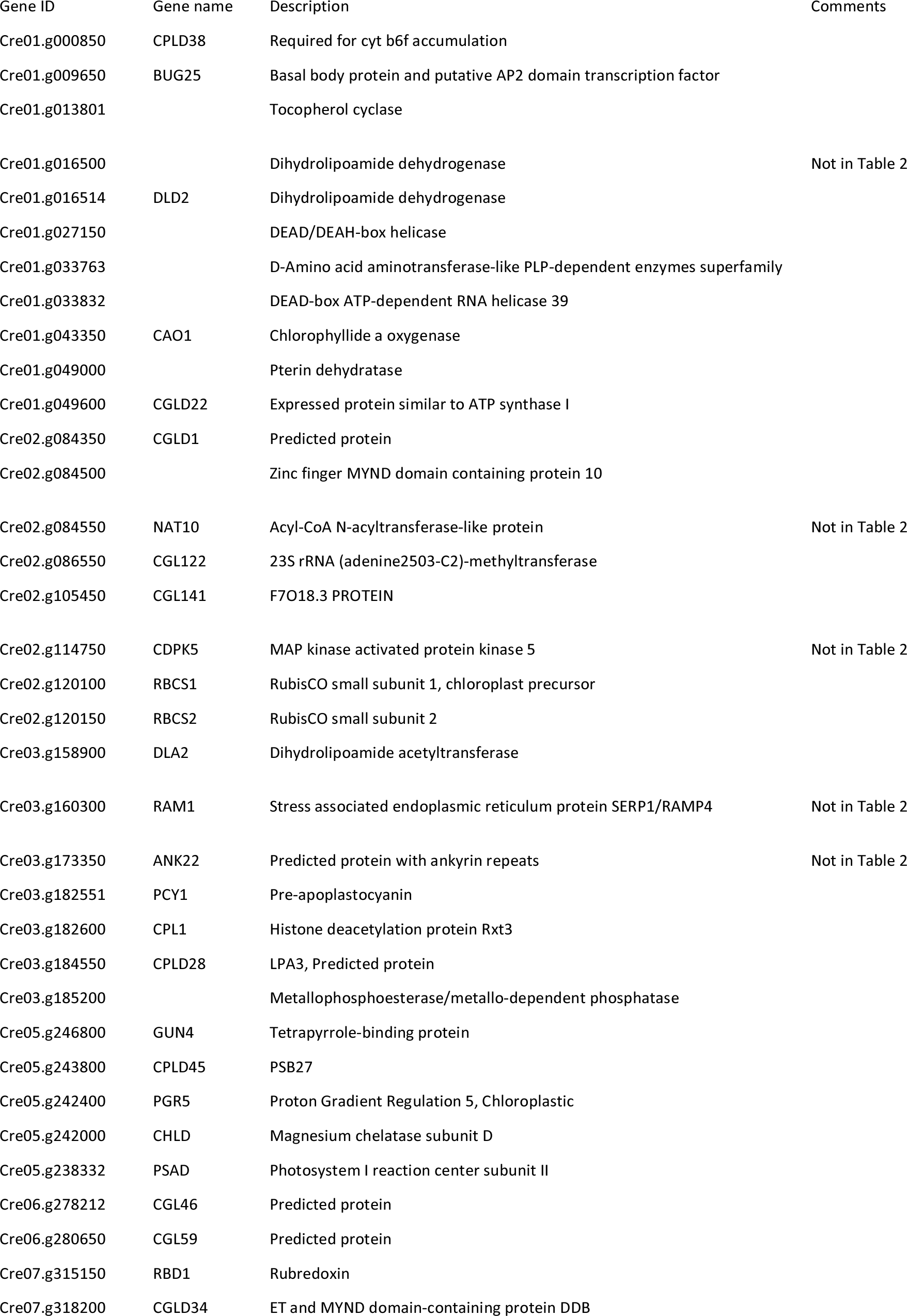
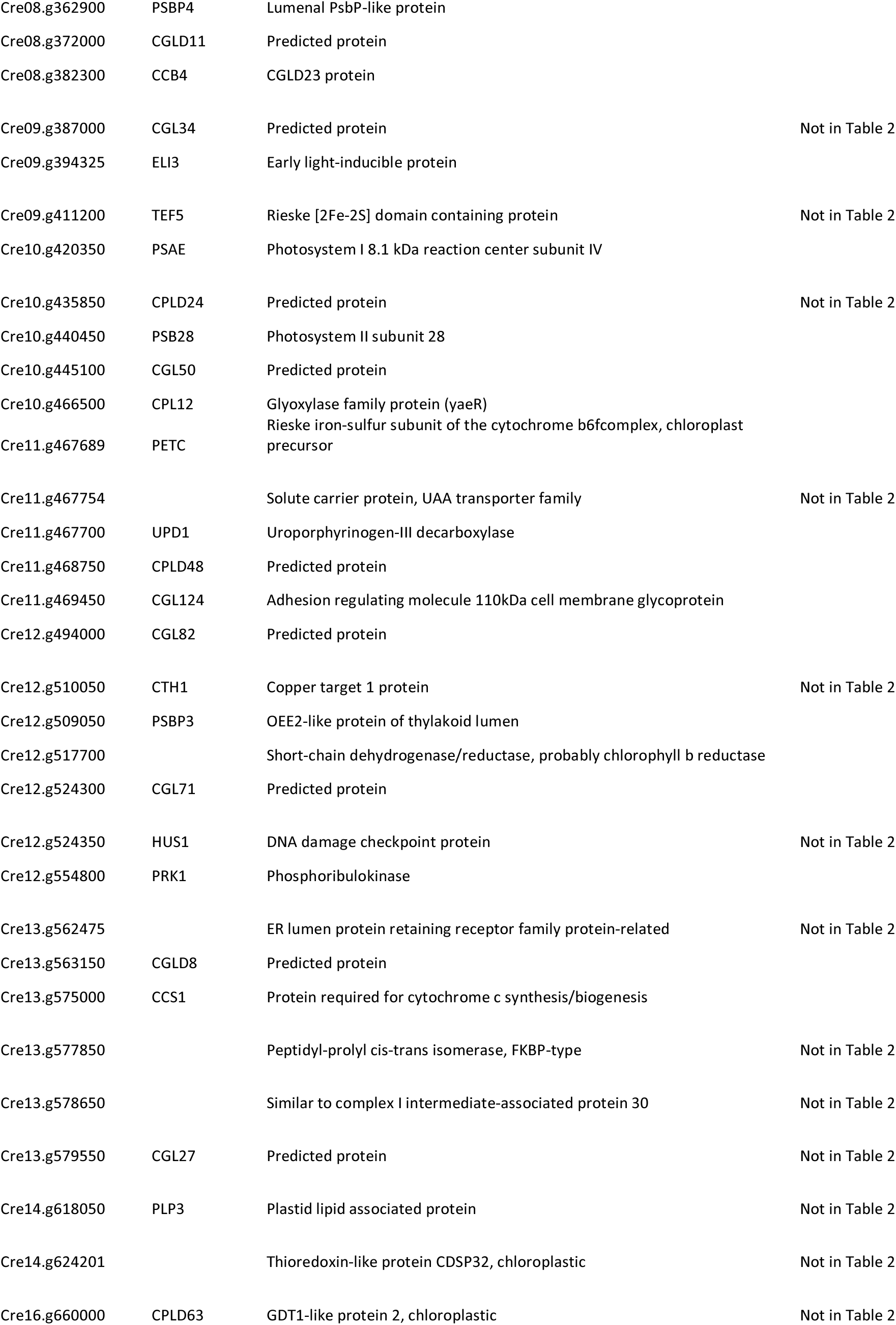
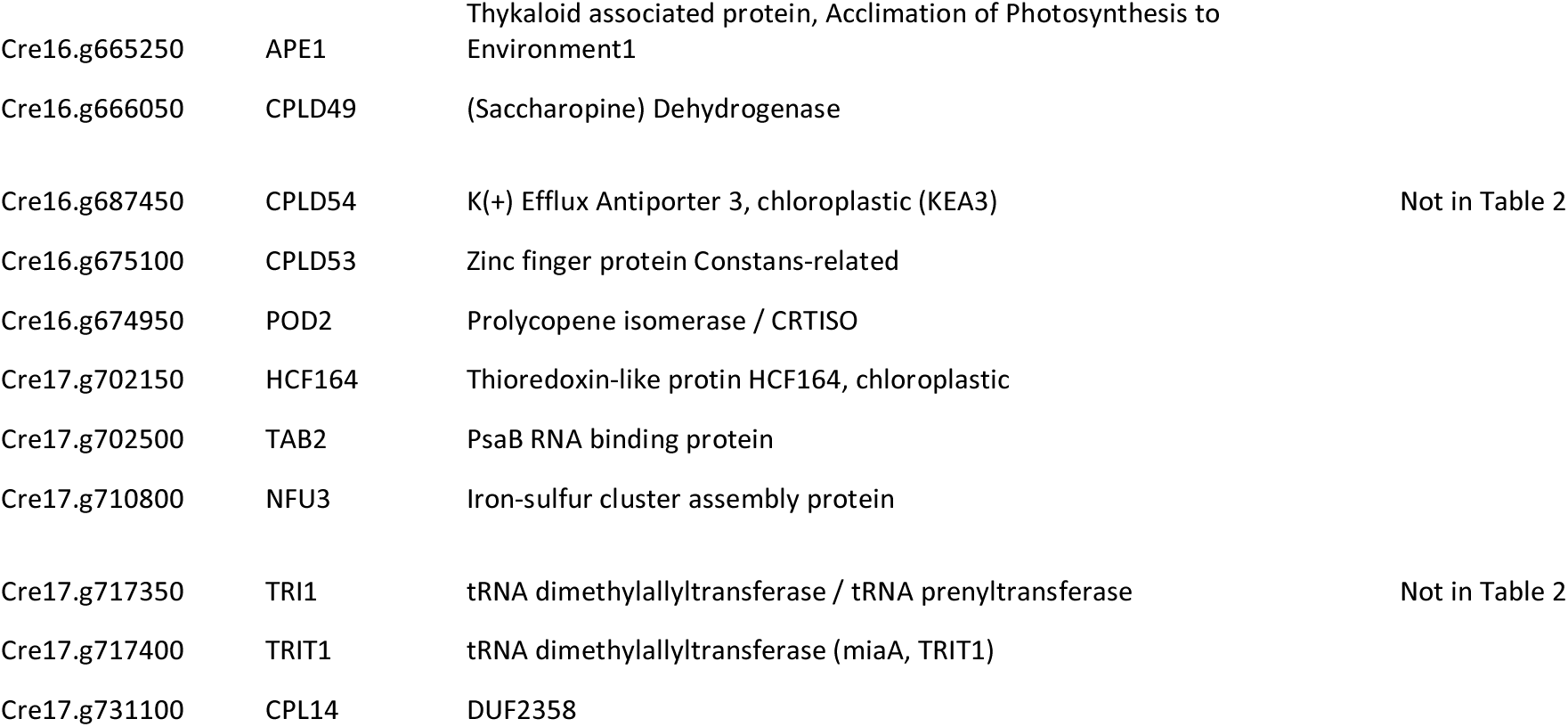
GreenCut2 genes affected in ARC.

| Gene ID | Gene name | Description | Comments |
| --- | --- | --- | --- |
| Cre01.g000850 | CPLD38 | Required for cyt b6f accumulation |  |
| Cre01.g009650 | BUG25 | Basal body protein and putative AP2 domain transcription factor |  |
| Cre01.g013801 |  | Tocopherol cyclase |  |
| Cre01.g016500 |  | Dihydrolipoamide dehydrogenase | Not in Table 2 |
| Cre01.g016514 | DLD2 | Dihydrolipoamide dehydrogenase |  |
| Cre01.g027150 |  | DEAD/DEAH-box helicase |  |
| Cre01.g033763 |  | D-Amino acid aminotransferase-like PLP-dependent enzymes superfamily |  |
| Cre01.g033832 |  | DEAD-box ATP-dependent RNA helicase 39 |  |
| Cre01.g043350 | CAO1 | Chlorophyllide a oxygenase |  |
| Cre01.g049000 |  | Pterin dehydratase |  |
| Cre01.g049600 | CGLD22 | Expressed protein similar to ATP synthase I |  |
| Cre02.g084350 | CGLD1 | Predicted protein |  |
| Cre02.g084500 |  | Zinc finger MYND domain containing protein 10 |  |
| Cre02.g084550 | NAT10 | Acyl-CoA N-acyltransferase-like protein | Not in Table 2 |
| Cre02.g086550 | CGL122 | 23S rRNA (adenine2503-C2)-methyltransferase |  |
| Cre02.g105450 | CGL141 | F7O18.3 PROTEIN |  |
| Cre02.g114750 | CDPK5 | MAP kinase activated protein kinase 5 | Not in Table 2 |
| Cre02.g120100 | RBCS1 | RubisCO small subunit 1, chloroplast precursor |  |
| Cre02.g120150 | RBCS2 | RubisCO small subunit 2 |  |
| Cre03.g158900 | DLA2 | Dihydrolipoamide acetyltransferase |  |
| Cre03.g160300 | RAM1 | Stress associated endoplasmic reticulum protein SERP1/RAMP4 | Not in Table 2 |
| Cre03.g173350 | ANK22 | Predicted protein with ankyrin repeats | Not in Table 2 |
| Cre03.g182551 | PCY1 | Pre-apoplastocyanin |  |
| Cre03.g182600 | CPL1 | Histone deacetylation protein Rxt3 |  |
| Cre03.g184550 | CPLD28 | LPA3, Predicted protein |  |
| Cre03.g185200 |  | Metallophosphoesterase/metallo-dependent phosphatase |  |
| Cre05.g246800 | GUN4 | Tetrapyrrole-binding protein |  |
| Cre05.g243800 | CPLD45 | PSB27 |  |
| Cre05.g242400 | PGR5 | Proton Gradient Regulation 5, Chloroplastic |  |
| Cre05.g242000 | CHLD | Magnesium chelatase subunit D |  |
| Cre05.g238332 | PSAD | Photosystem I reaction center subunit II |  |
| Cre06.g278212 | CGL46 | Predicted protein |  |
| Cre06.g280650 | CGL59 | Predicted protein |  |
| Cre07.g315150 | RBD1 | Rubredoxin |  |
| Cre07.g318200 | CGLD34 | ET and MYND domain-containing protein DDB |  |
| Cre08.g362900 | PSBP4 | Lumenal PsbP-like protein |  |
| Cre08.g372000 | CGLD11 | Predicted protein |  |
| Cre08.g382300 | CCB4 | CGLD23 protein |  |
| Cre09.g387000 | CGL34 | Predicted protein | Not in Table 2 |
| Cre09.g394325 | ELI3 | Early light-inducible protein |  |
| Cre09.g411200 | TEF5 | Rieske [2Fe-2S] domain containing protein | Not in Table 2 |
| Cre10.g420350 | PSAE | Photosystem I 8.1 kDa reaction center subunit IV |  |
| Cre10.g435850 | CPLD24 | Predicted protein | Not in Table 2 |
| Cre10.g440450 | PSB28 | Photosystem II subunit 28 |  |
| Cre10.g445100 | CGL50 | Predicted protein |  |
| Cre10.g466500 | CPL12 | Glyoxylase family protein (yaeR) |  |
| Cre11.g467689 | PETC | Rieske iron-sulfur subunit of the cytochrome b6f complex, chloroplast precursor |  |
| Cre11.g467754 |  | Solute carrier protein, UAA transporter family | Not in Table 2 |
| Cre11.g467700 | UPD1 | Uroporphyrinogen-III decarboxylase |  |
| Cre11.g468750 | CPLD48 | Predicted protein |  |
| Cre11.g469450 | CGL124 | Adhesion regulating molecule 110kDa cell membrane glycoprotein |  |
| Cre12.g494000 | CGL82 | Predicted protein |  |
| Cre12.g510050 | CTH1 | Copper target 1 protein | Not in Table 2 |
| Cre12.g509050 | PSBP3 | OEE2-like protein of thylakoid lumen |  |
| Cre12.g517700 |  | Short-chain dehydrogenase/reductase, probably chlorophyll b reductase |  |
| Cre12.g524300 | CGL71 | Predicted protein |  |
| Cre12.g524350 | HUS1 | DNA damage checkpoint protein | Not in Table 2 |
| Cre12.g554800 | PRK1 | Phosphoribulokinase |  |
| Cre13.g562475 |  | ER lumen protein retaining receptor family protein-related | Not in Table 2 |
| Cre13.g563150 | CGLD8 | Predicted protein |  |
| Cre13.g575000 | CCS1 | Protein required for cytochrome c synthesis/biogenesis |  |
| Cre13.g577850 |  | Peptidyl-prolyl cis-trans isomerase, FKBP-type | Not in Table 2 |
| Cre13.g578650 |  | Similar to complex I intermediate-associated protein 30 | Not in Table 2 |
| Cre13.g579550 | CGL27 | Predicted protein | Not in Table 2 |
| Cre14.g618050 | PLP3 | Plastid lipid associated protein | Not in Table 2 |
| Cre14.g624201 |  | Thioredoxin-like protein CDSP32, chloroplastic | Not in Table 2 |
| Cre16.g660000 | CPLD63 | GDT1-like protein 2, chloroplastic | Not in Table 2 |
| Cre16.g665250 | APE1 | Thylakoid associated protein, Acclimation of Photosynthesis to Environment1 |  |
| Cre16.g666050 | CPLD49 | (Saccharopine) Dehydrogenase |  |
| Cre16.g687450 | CPLD54 | K(+) Efflux Antiporter 3, chloroplastic (KEA3) | Not in Table 2 |
| Cre16.g675100 | CPLD53 | Zinc finger protein Constans-related |  |
| Cre16.g674950 | POD2 | Prolycopene isomerase / CRTISO |  |
| Cre17.g702150 | HCF164 | Thioredoxin-like protein HCF164, chloroplastic |  |
| Cre17.g702500 | TAB2 | PsaB RNA binding protein |  |
| Cre17.g710800 | NFU3 | Iron-sulfur cluster assembly protein |  |
| Cre17.g717350 | TRI1 | tRNA dimethylallyltransferase / tRNA prenyltransferase | Not in Table 2 |
| Cre17.g717400 | TRIT1 | tRNA dimethylallyltransferase (miaA, TRIT1) |  |
| Cre17.g731100 | CPL14 | DUF2358 |  |

We were able to identify a higher-confidence candidate gene for 348 out of 436 mutants with simple insertions. Because there were multiple alleles of 59 genes, this resulted in 273 higher-confidence candidate genes, which are shown in Table 2 (and S4 Table with additional details and references). This list includes genes known to be important for photosynthesis, photoprotection, and peripheral functions (S4 Table, Column “Inferred function from Cr and other photosynthetic organisms”). 106 gene products were predicted to be targeted to the chloroplast by protein targeting software Predalgo (https://giavap-genomes.ibpc.fr/cgi-bin/predalgodb.perl?page=main) (44), and among those, 61 were also predicted to be targeted to plastids by ChloroP (http://www.cbs.dtu.dk/services/ChloroP/) (45) (Table 2, S4 Table). 55 GreenCut2 genes are within this higher-confidence list, leaving 23 GreenCut2 genes that were not chosen because there was a stronger candidate gene (see column “Comments” in Table 2), an indication that not all GreenCut2 genes may be critical for photosynthesis. Among the 273 candidates, the photosynthetic functions of 68 genes have been previously described in *Chlamydomonas*, land plants, or cyanobacteria. This leaves 205 genes whose functions remain to be studied in context of photosynthesis, 47 of which have no annotation (S4 Table).

**Table 2.**
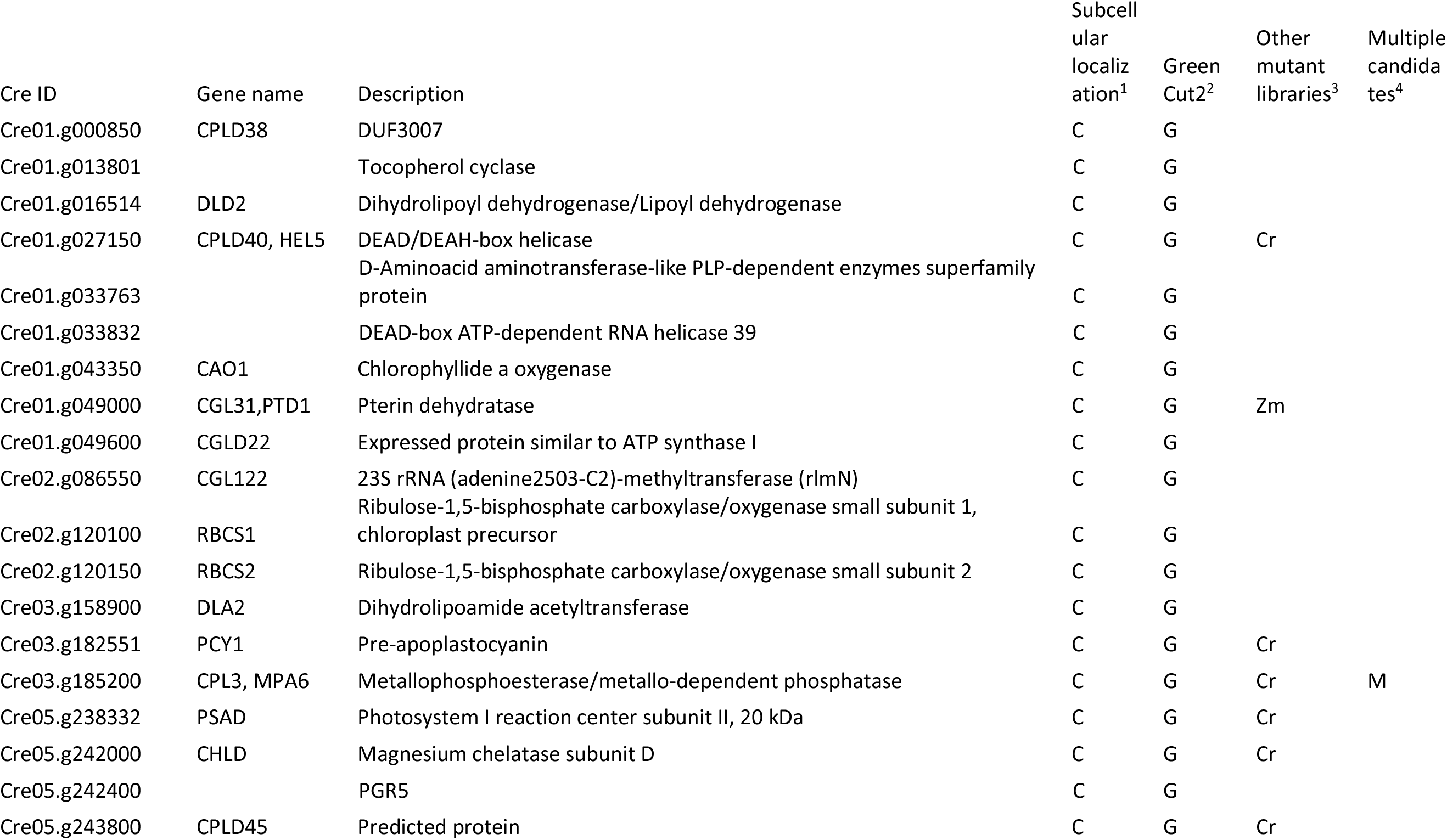
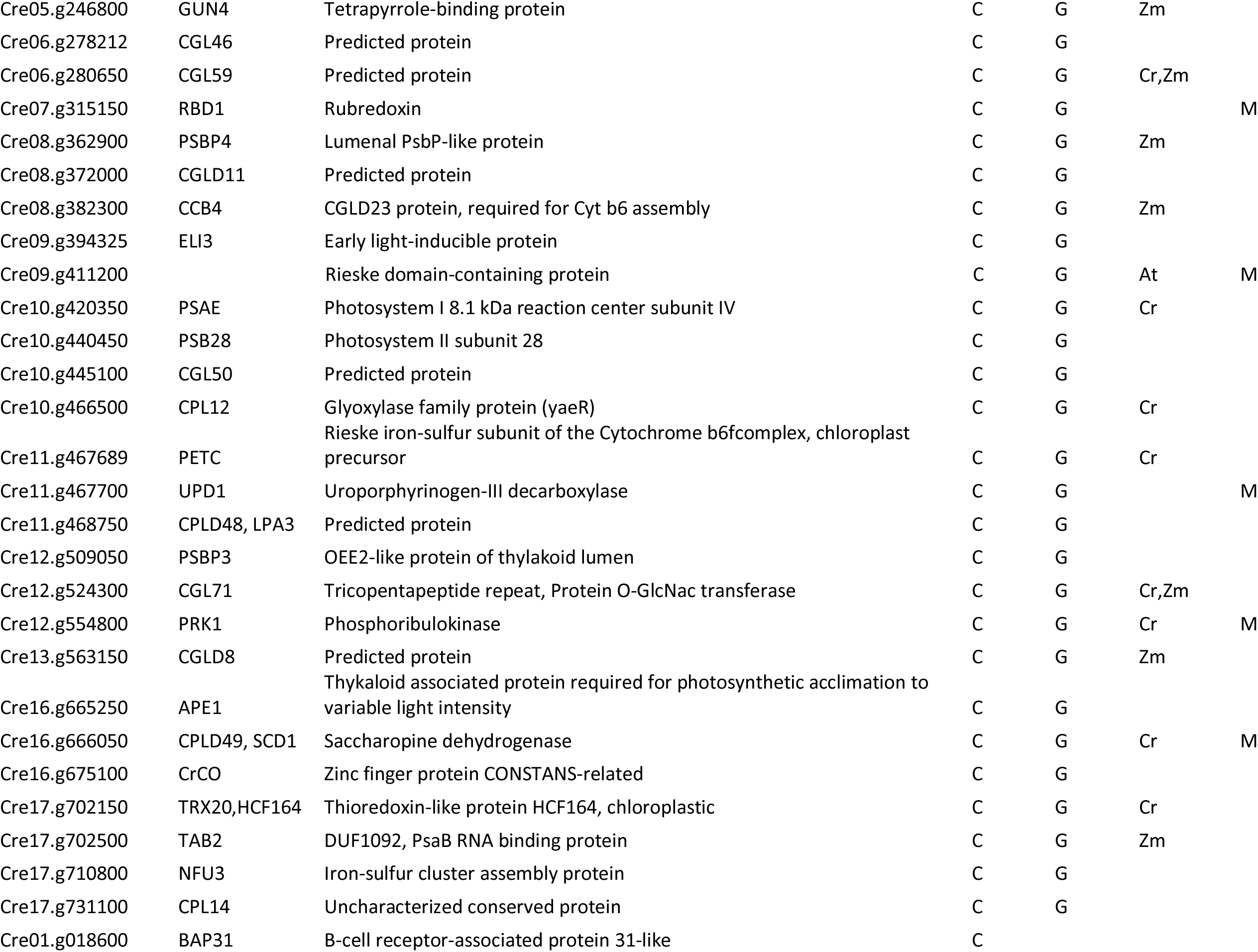
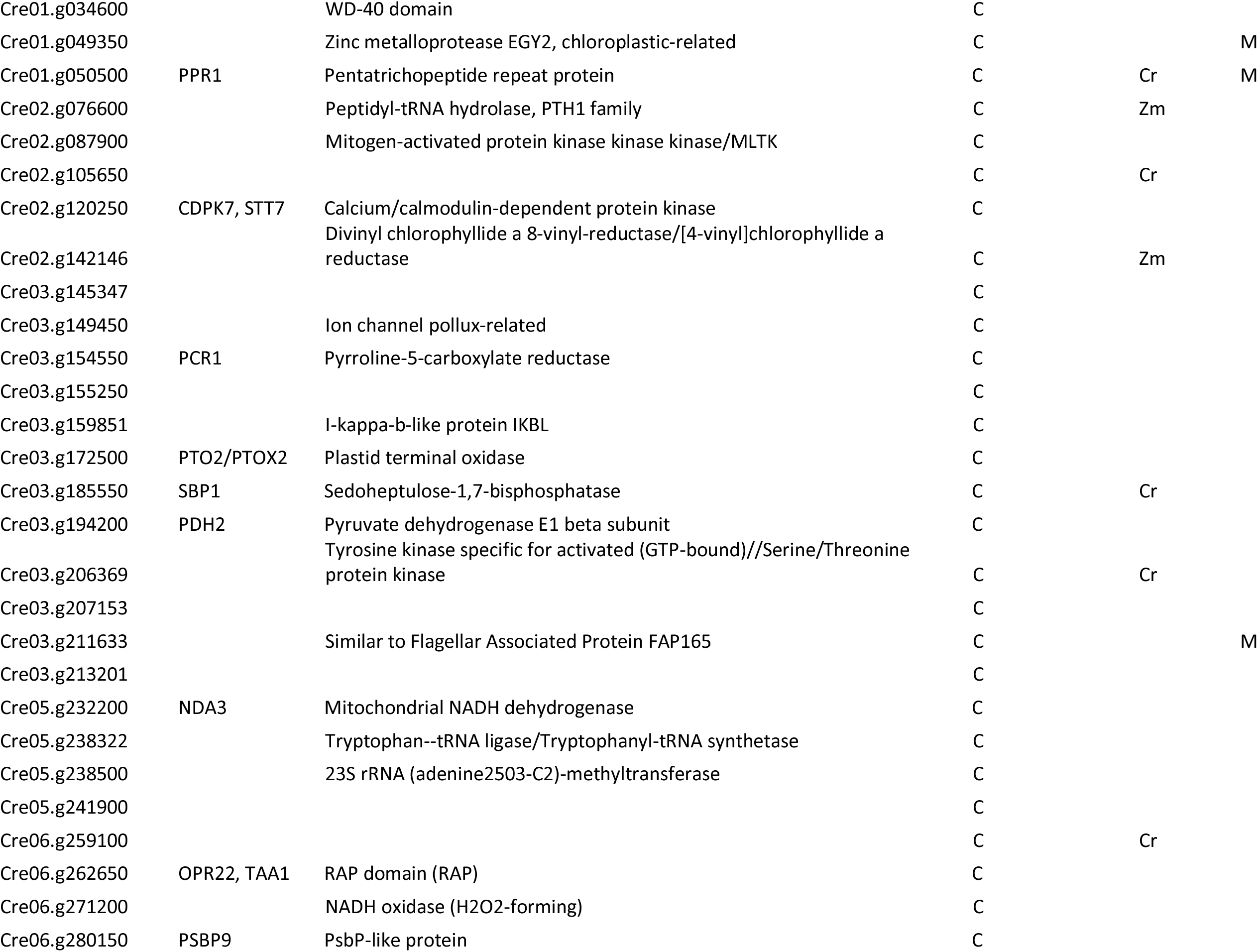
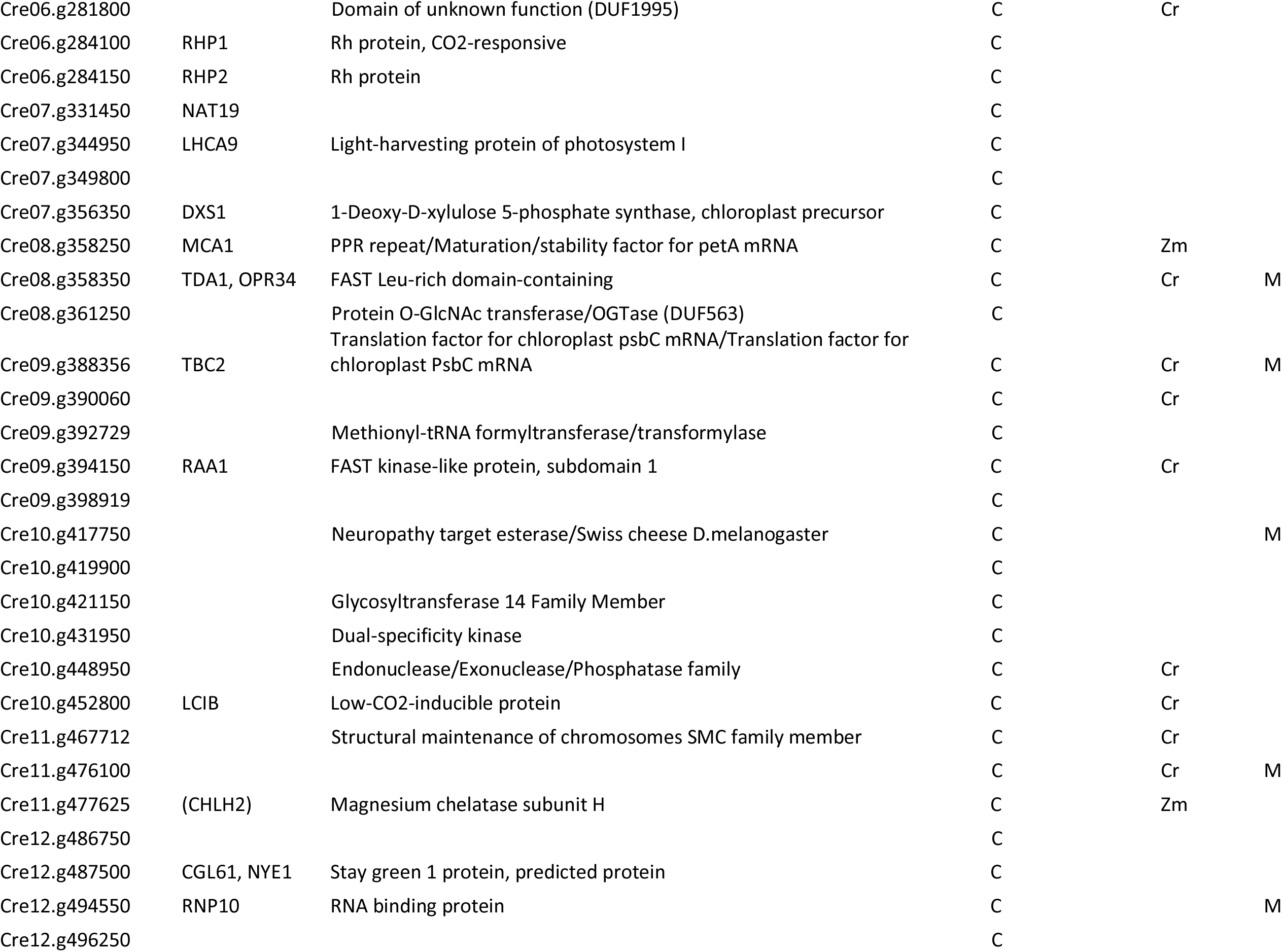
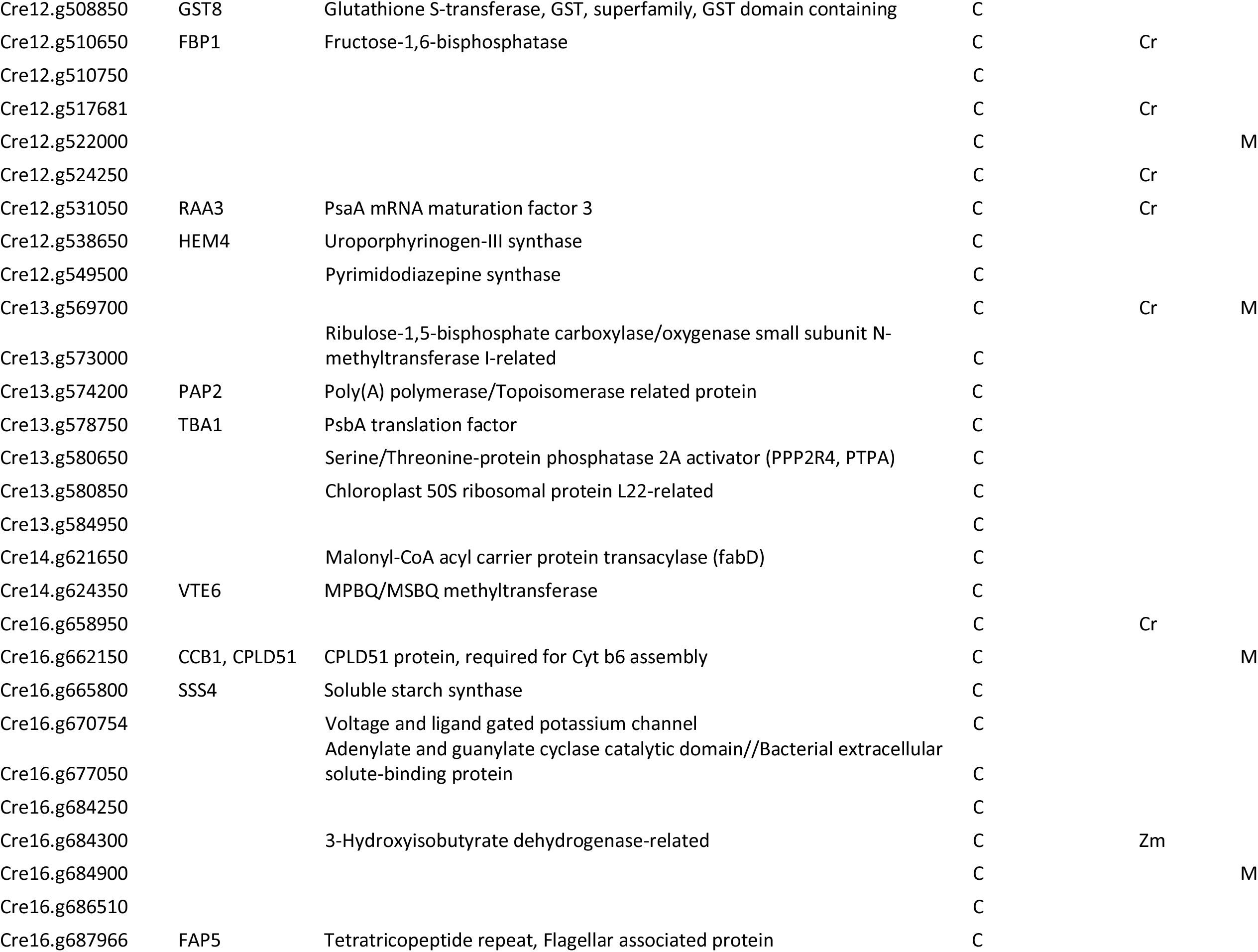
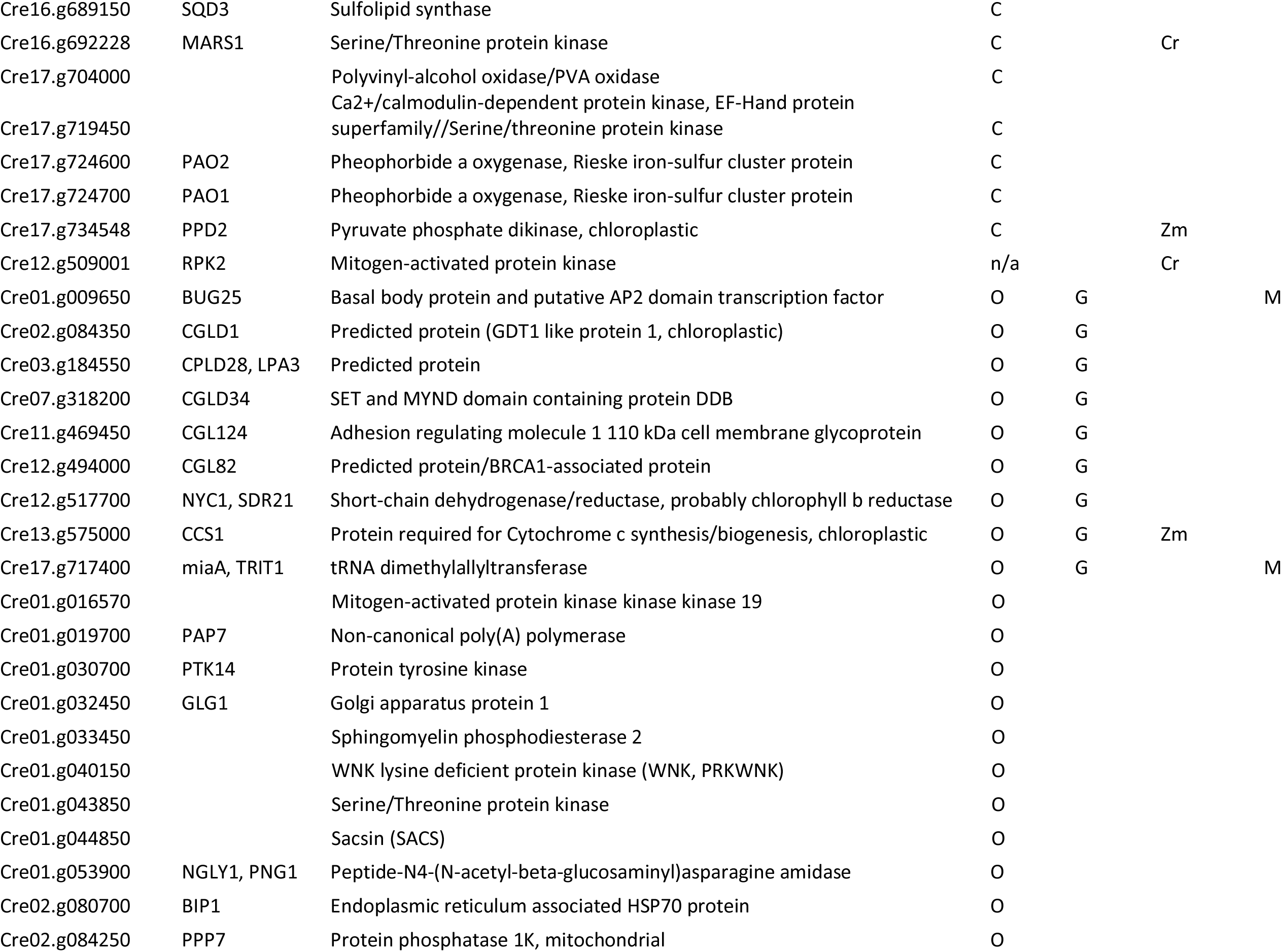
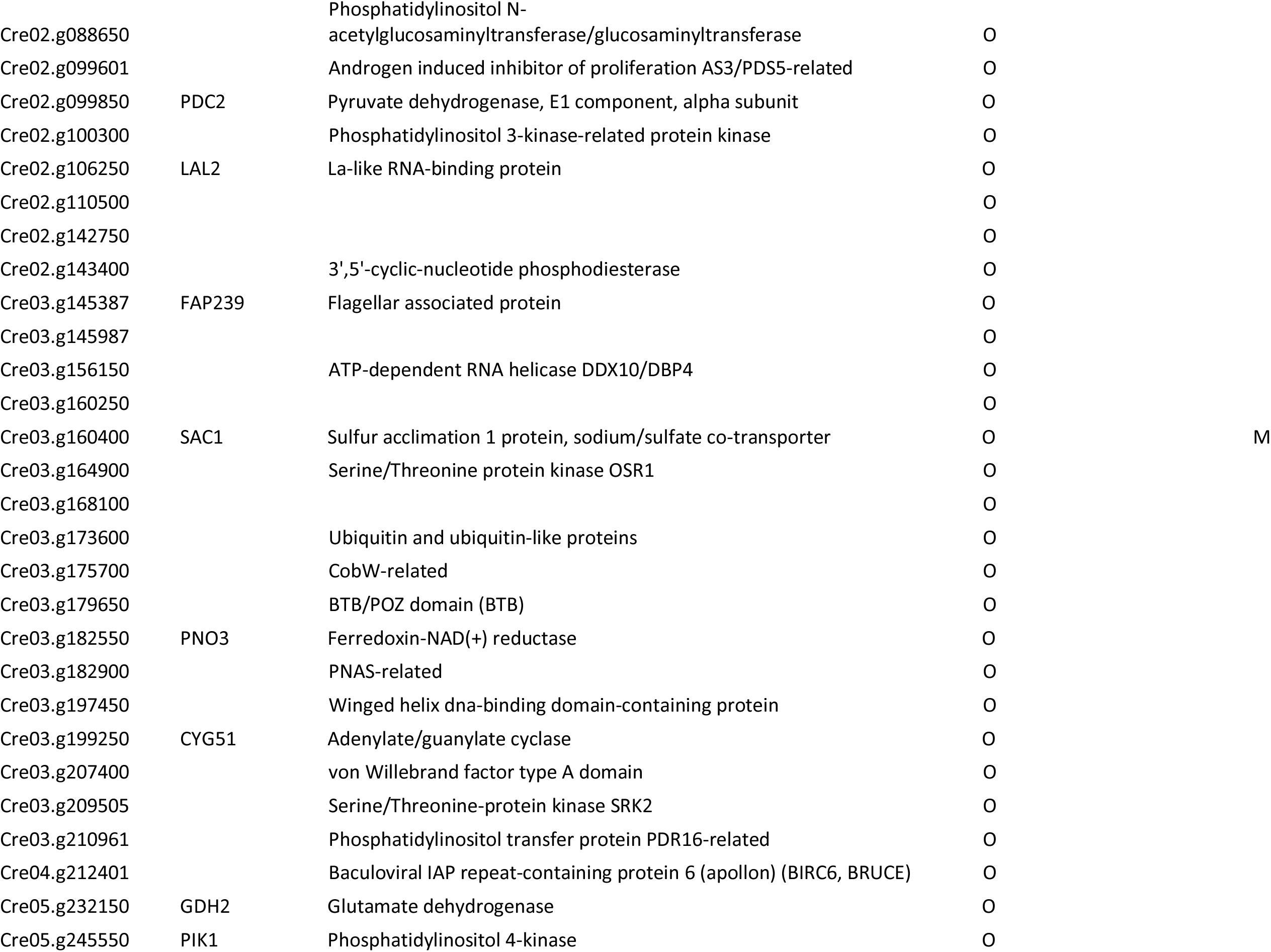
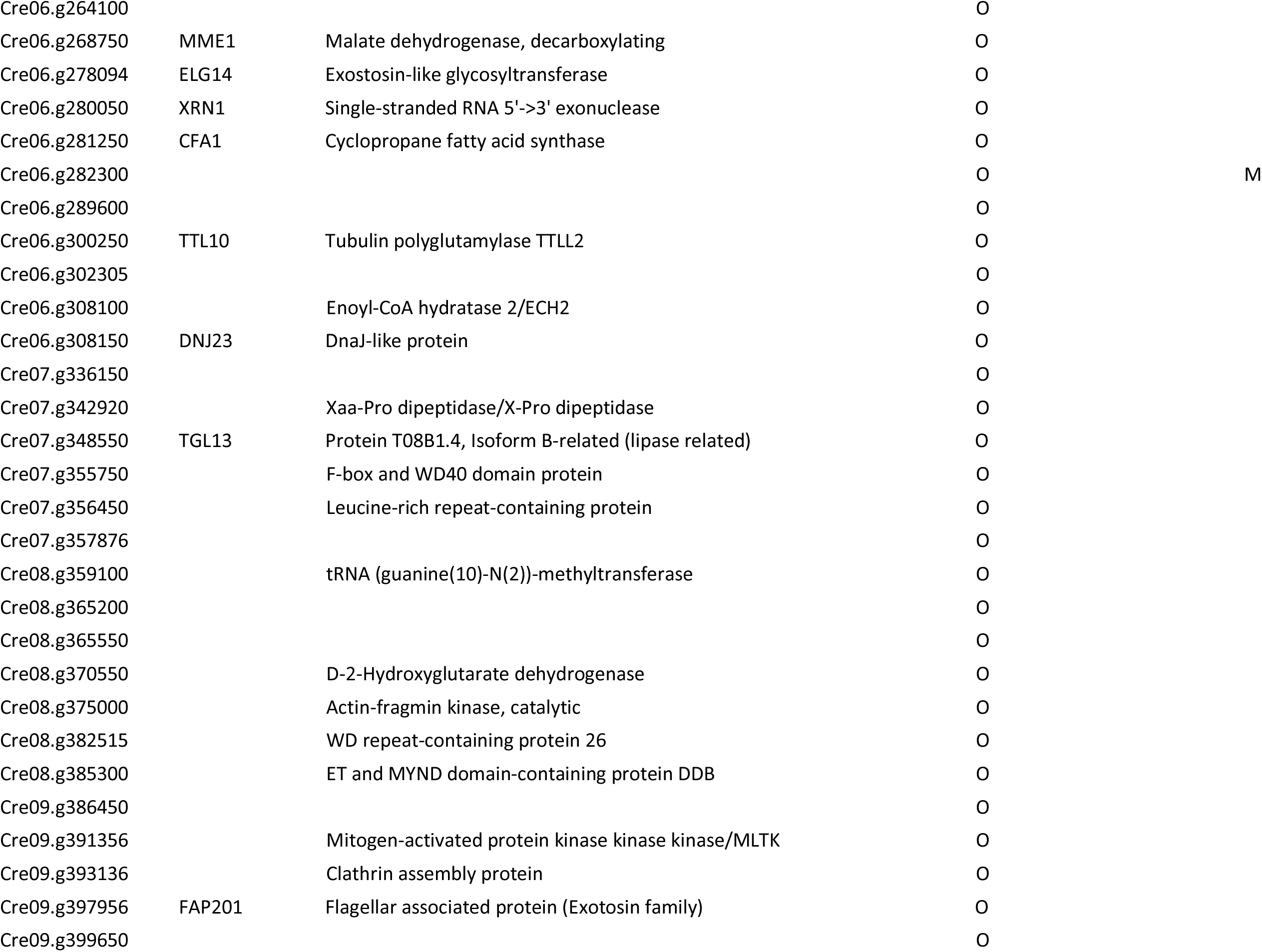
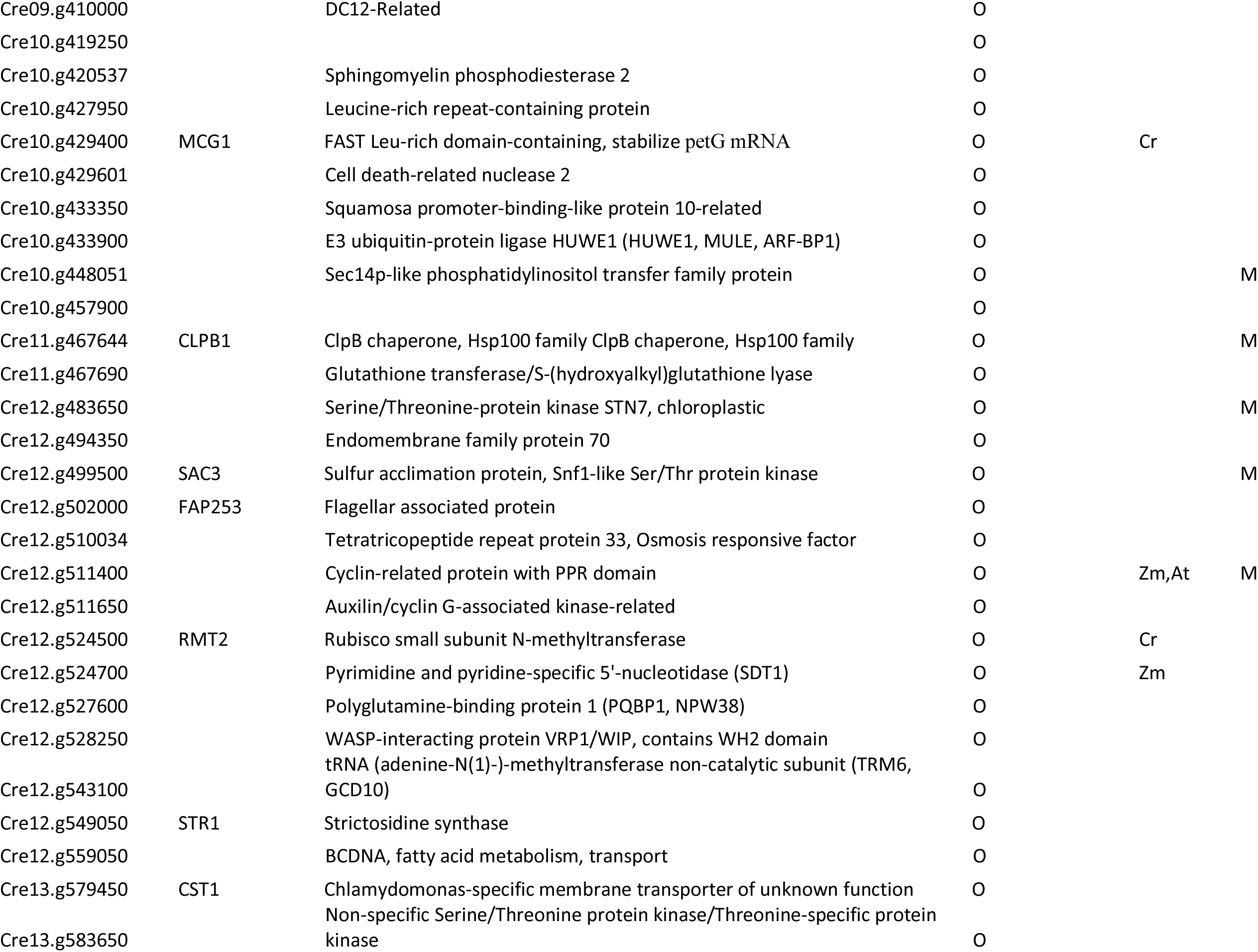
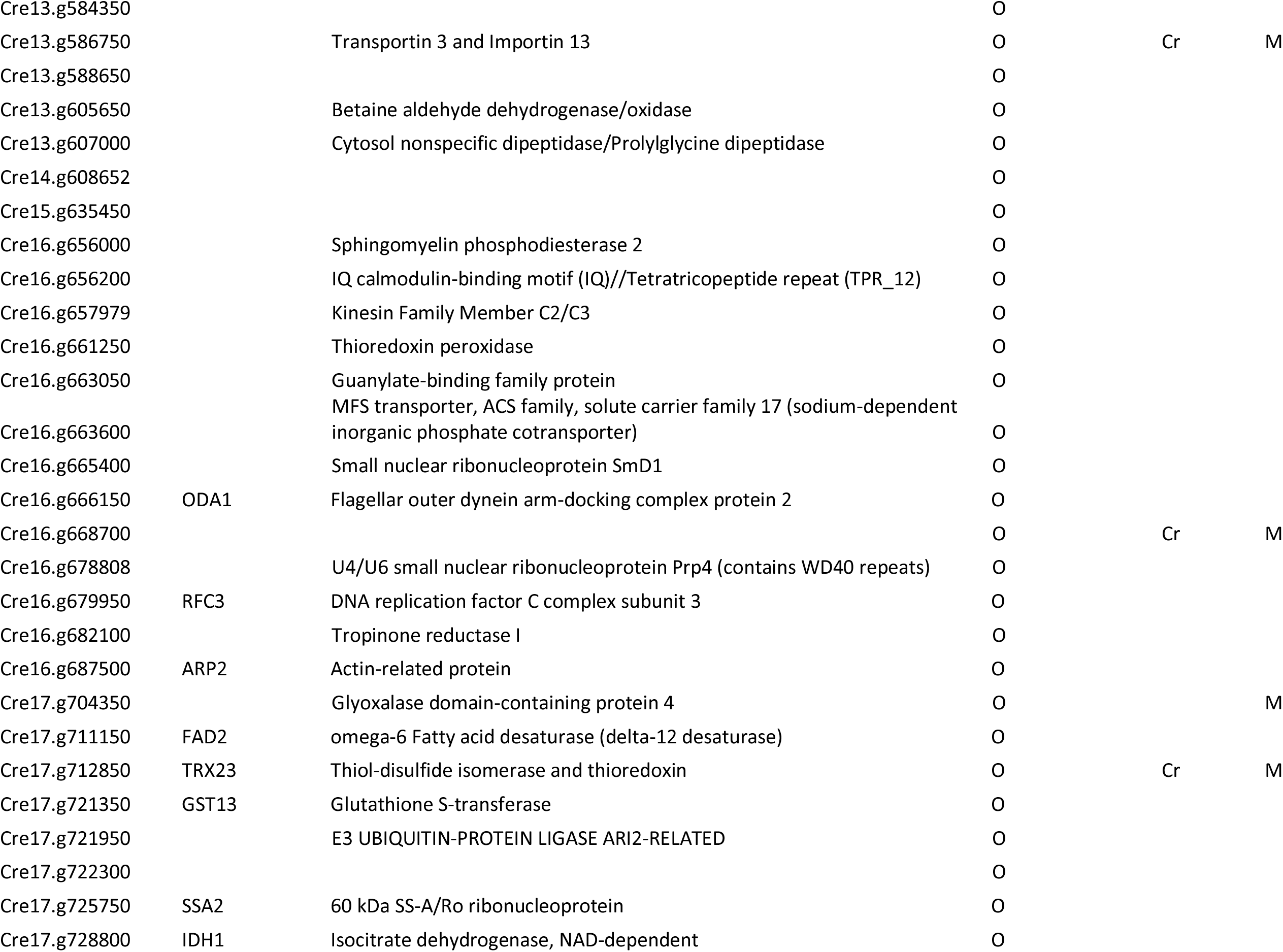
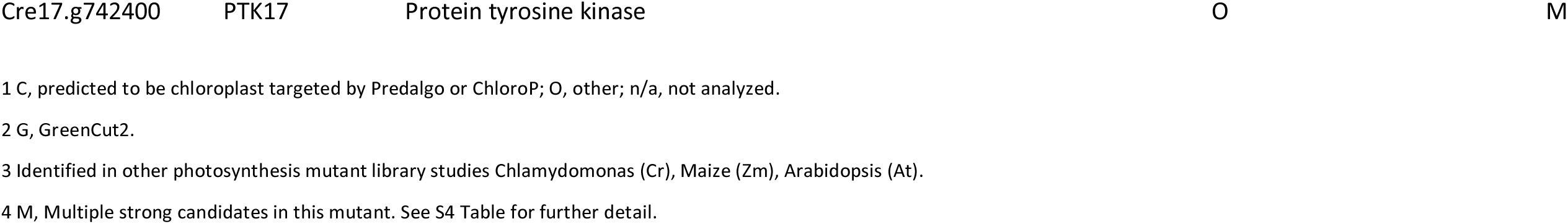
Higher confidence photosynthesis candidate genes.

## Discussion

We successfully used high-throughput, low-coverage WGS for the identification of plasmid insertion sites in our *Chlamydomonas* photosynthesis mutant collection (ARC). This approach has a much higher efficiency than PCR-based FST isolation. From the larger collection of 2800 mutants (7) from which ARC was derived, we recovered FSTs from only 17% of the mutants, whereas our WGS identified insertions in 509 out of 581 non-redundant ARC mutants (88% success among the population). We attribute this improvement to the fact that insertion site identification by WGS is not dependent on the intactness or sequence continuity of the inserted plasmid sequence, and therefore WGS overcomes complications such as plasmid concatemerization and loss of plasmid ends to which PCR primers need to anneal. Most importantly, it completely bypasses the need for PCR from the GC- and repeat-rich genome of *Chlamydomonas*. Even with relatively low average WGS coverage (∼7x), we also identified 68 deletions that were not associated with plasmid insertions, some of which may be causative mutations for photosynthesis-related phenotypes that are unlinked to the plasmid insertion in specific mutants.

A previous study using WGS to identify DNA insertion events in *Chlamydomonas* (21) provides the most direct comparison with our results. Lin et al. (2018) analyzed paromomycin-resistant insertional mutants derived from electroporation instead of the glass bead transformation method that we used to generate either paromomycin- or zeocin-resistant mutants (9). They sequenced 20 transformants in 10 pools of two strains and verified 38 insertions, obtaining an average of 1.9 insertions per strain. In contrast, we found a total of 554 insertions in 509 mutants, resulting in a lower average of ∼1.1 insertions per mutant. Lin et al. (2018) found that more than half (11 of 20) of their strains had more than one insertion event, and a larger collection of 1935 mutants derived from electroporation exhibited multiple insertions in 26% of strains (10). We found multiple insertions in 8% (43 out of 509) of the ARC mutants, suggesting that glass bead transformation of *Chlamydomonas* results in a higher frequency of single-copy insertions. Lin et al. (2018) identified one-sided insertions in ∼40% of their mutants, whereas we observed only ∼4% (21 out of 554 insertion events), despite the lower average WGS coverage in our study (∼7x vs. ∼15x). The frequency of complex rearrangements in our study (19%) was comparable to that observed by Lin et al. (25%), however, as previously noted by us and others (7,10,21,46), glass bead transformation seems to be frequently associated with larger deletions of genomic DNA at the sites of DNA insertion than electroporation, a finding that was clearly evident in our WGS data (Figure 3A).

In part because of the occurrence of larger deletions, 1405 genes were disrupted in 509 ARC mutants. As expected, this list is enriched for genes that encode proteins with annotated functions in photosynthesis and tetrapyrrole synthesis, and it includes 78 GreenCut2 genes (31). We examined the affected genes in each mutant to identify possible causative genes using several criteria, including GreenCut2 membership, existence of protein domains suggestive of a function in photosynthesis, and occurrence of multiple mutant alleles in the ARC. We also searched for overlaps with available photosynthesis mutant datasets, namely CLiP (*Chlamydomonas*), PML (maize), DEPI (*Arabidopsis*), and those found co-expressed with photosynthesis genes (*Chlamydomonas*). The CLiP collection has been used to identify mutants that are defective in photosynthetic growth in pooled cultures (9). This study identified 303 candidate photosynthesis genes. We identified 41 of those 303 genes in our list of 273 higher-confidence genes (Table 2, S4 Table). This overlap is lower than might be expected but could be explained simply by the fact that both the CLiP and ARC mutant collections are based on a total of ∼60,000 insertional mutants, which is not sufficient to saturate the *Chlamydomonas* genome for mutations affecting photosynthesis. The maize PML consists of approximately 2100 photosynthesis mutants that contain 50 to 100 *Mu* transposable elements per individual. It is estimated to be a saturated collection with 3-4 mutant alleles for ∼600 genes (42). The FSTs of this library were obtained with Illumina sequencing of fragmented gDNA that was enriched for the *Mu* element (22). Our higher-confidence candidate gene list overlapped with 17 genes identified from the maize PML (http://pml.uoregon.edu/photosyntheticml.html). DEPI screening of 300 *Arabidopsis* mutants affecting genes that encode chloroplast-targeted proteins (Chloroplast 2010 project, http://www.plastid.msu.edu/) identified 12 mutants with altered photosynthetic response (43). These mutants likely represent disruption in genes that are conditionally important in acclimation to changing light environments. Two of the 12 genes found through DEPI overlapped with our higher-confidence photosynthesis candidate gene list. The largest overlap (84 genes) was observed between our higher-confidence list and the group of photosynthesis-related genes defined based on co-expression analysis (47).

For two of the higher-confidence photosynthesis genes, *LPA3* and *PSBP4*, we validated the insertion-associated lesions for four of the ARC mutants and demonstrated their requirement for photoautotrophic growth (Figure 5). LPA3 is a GreenCut2 protein (CPLD28) that contains a DUF1995 domain. Insertion mutants containing large or small deletions in *LPA3* (Cre03.g184550) were acetate-requiring and exhibited a severe defect in PSII function even in the dark, as evidenced by F_v_/F_m_ values near zero (Figure 5). Mutants affecting Cre02.g105650 and Cre10.g441650, two *Chlamydomonas* genes coding for proteins similar to *Arabidopsis* LPA2, were not found in the ARC. However, there are two additional genes encoding DUF1995 proteins in the *Chlamydomonas* genome, Cre06.g281800 and Cre08.g369000. The mutant CAL038_02_36 is disrupted in Cre06.g281800. It does not grow photoautotrophically but is able to grow in LL and HL in the presence of acetate. Interestingly, this mutant also has an F_v_/F_m_ of zero in the dark (S1 Table). The severe phenotypes of these mutants in *Chlamydomonas* indicate non-overlapping functions in PSII assembly of the gene products of *LPA3* and Cre06.g281800.

PSBP (encoded by *PSBP1*/*OEE2* in *Chlamydomonas*) together with PSBO and PSBQ constitute the oxygen-evolving complex (OEC) of PSII (48, 49). In green algae and plants, PSBP appears to have expanded into a large family of proteins sharing similar domains beyond the canonical PSBP of the OEC. The *Chlamydomonas* genome contains 13 additional genes encoding proteins with PsbP-like domains whose individual functions are unknown. We showed that PSBP4 is required for photoautotrophic growth in *Chlamydomonas*, ruling out redundancy in its function with other PSBP-like domain-containing proteins. An *Arabidopsis* ortholog of CrPSBP4 (AT4g15510, PPD1) has been shown to play a role in PSI assembly (40, 41), which is consistent with the light-sensitivity of our *psbp4-1* mutant. Two other members of the PSBP family, *PSBP3*, and *PSBP9*, were found to be disrupted in the ARC. The large family of PSBP-like domain-containing proteins is speculated to have resulted in divergence of their functions (50), and the availability of mutants in these genes should help to reveal their functions.

Of the 273 higher-confidence candidate photosynthesis genes that we curated based on WGS analysis of the ARC, only 68 have a previously demonstrated function in photosynthesis. This is similar to the results of pooled growth analysis of ∼60,000 *Chlamydomonas* insertional mutants by Li et al. (2019), which revealed 303 candidate photosynthesis genes, of which only 65 have previously known roles in photosynthesis (9). Thus, 238 genes in the study of Li et al. (2019) and 205 genes in our study remain to be analyzed experimentally to determine their specific functions in photosynthesis. Moreover, the fact that only 42 genes are shared by these two sets of candidate photosynthesis genes suggests that there are still many more photosynthesis genes that remain to be identified, which highlights the enormous potential for future validation and discovery of new proteins involved in oxygenic photosynthesis.

## Material and methods

### Strains and culture conditions

Mutants described in this work were generated from wild-type strain 4A+ (CC-4051 in the 137c background. Cells were grown mixotrophically (ac) on Tris-acetate-phosphate (TAP) medium and photoautotrophically (min) on minimal high-salt medium (HS) medium (51) in low light (LL) of 60-80 μmol photons m^-2^ s^-1^ and high light (HL) of 350-400 μmol photons m^-2^ s^-1^. LL and HL conditions were obtained using GE F25T8/SPX41/ECO and Sylvania F72T12/CW/VHO fluorescent bulbs, respectively.

### Genomic DNA preparation and whole-genome sequencing

*Chlamydomonas* cultures were grown in 20 mL TAP to stationary phase, and genomic DNA was extracted using an alkaline lysis buffer (50 mM Tris-HCl (pH 8), 200 mM NaCl, 20 mM EDTA, 2% SDS, 1% PVP 40,000, 1 mg/mL Proteinase K) followed by phenol-chloroform extraction. DNA was collected, washed and eluted using DNeasy Plant mini-columns (QIAGEN). The resulting quality of the DNA was confirmed to be A_260_/A_280_ of approximately 1.8 and A_260_/A_230_ of >2. Plate-based DNA library preparation for Illumina sequencing was performed on the PerkinElmer Sciclone NGS robotic liquid handling system using Kapa Biosystems library preparation kit. 200 ng of sample DNA was sheared to 600 bp using a Covaris LE220 focused ultrasonicator. The sheared DNA fragments were size selected by double-SPRI, and then the selected fragments were end-repaired, A-tailed, and ligated with Illumina-compatible sequencing adaptors from IDT containing a unique molecular index barcode for each sample library. The prepared libraries were quantified using KAPA Biosystem’s next-generation sequencing library qPCR kit and run on a Roche LightCycler 480 real-time PCR instrument. The quantified libraries were then multiplexed with other libraries, and the pool of libraries was then prepared for sequencing on the Illumina HiSeq sequencing platform utilizing a TruSeq paired-end cluster kit, v4, and Illumina’s cBot instrument to generate a clustered flow cell for sequencing. Sequencing of the flow cell was performed on the Illumina HiSeq2500 sequencer using HiSeq TruSeq SBS sequencing kits, v4, following a 2×150 indexed run recipe. The reads were aligned to the reference genome using BWA-mem. To identify plasmid insertion sites, discordant paired-end reads with one end mapping to the plasmid used for mutagenesis and the other to a chromosome location were mapped and manually validated for each mutant using Integrated Genome Viewer (IGV) (http://software.broadinstitute.org/software/igv/home). Putative structural variations unpaired to the plasmid sequence were called using a combination of BreakDancer (filtered to quality 90+) and Pindel and manually validated using IGV. Resulting genome sequences of 79 mutants were not unique (33 were duplicated, three were triplicated and one was quadruplicated). In all cases the mutants sharing similar sequences came from the same agar plate and sequencing plate, suggesting that it could be due to an error at the genome extraction step or in maintenance of the mutant strains; these mutants were not included in further analysis.

### Molecular analyses of mutants by PCR and mutant complementation

Deletions predicted from genome sequences were confirmed by using PCR primers that anneal proximal to the borders and within the deletions. The insertion of the plasmid sequence accompanied by a 4 bp-deletion in *lpa3-3* was sequenced from the PCR product from the predicted region. Primers used for PCRs indicated in Figure 4 are listed in Supplemental S4 Table. For complementation of *lpa3-1*, *lpa3-2*, and *lpa3-3*, a 3531 bp genomic fragment containing the full length *CrLPA3* gene (Cre03.g184550) with 1209 bp upstream of the start codon and 719 bp downstream of the stop codon was amplified using primers Comp11F and Comp11R. This fragment was subsequently Gibson cloned into the vector pSP124S using primers PS1362 and PS1363 to inverse PCR around pSP124S. For complementation of mutant *psbp4-1*, a 3246 bp genomic fragment containing the full length *CrPSBP4* gene (Cre08.g362900), including 1209 bp upstream of the start codon and 719 bp downstream of the stop codon, was amplified using primers Comp12F and Comp12R and similarly cloned into vector pSP124S. Primer sequences are listed in supplemental S4 Table. Constructs for complementation were transformed into the respective mutants using the glass bead method (52). Colonies were selected on 10 μM zeocin TAP agar plates and screened for rescued individuals by measuring F_v_/F_m_ as described below.

### F_v_/F_m_ measurement

*Chlamydomonas* strains were grown on agar plates in Dark+ac, LL-min, or HL-min, and F_v_/F_m_ (F_m_-F_o_/F_m_) was measured using a chlorophyll fluorescence video imager (IMAG-MAX/L, WALZ). Plates with the streaks of strains were dark-acclimated for 30 min and exposed to a pulse of saturating light (4000 μmol photons m^-2^ s^-1^). Fluorescence images of F_m_ and F_o_ were captured during saturating pulses, and false-color images of F_v_/F_m_ were generated.

## Supporting information

S1 Appendix Table S4 reference list

## Acknowledgments

We thank Alice Barkan for sharing the data for PML to compare with ARC higher-confidence candidate genes and Sabeeha Merchant, Masakazu Iwai, and Dhruv Patel for critical reading of the manuscript. This work was supported by the U.S. Department of Energy, Office of Science, Basic Energy Sciences, Chemical Sciences, Geosciences, and Biosciences Division under field work proposal 449B. The work conducted by the U.S. Department of Energy Joint Genome Institute, a DOE Office of Science User Facility, is supported by the Office of Science of the U.S. Department of Energy under Contract No. DE-AC02-05CH11231. K.K.N. is an investigator of the Howard Hughes Medical Institute.

## Supporting information

S1 Table. Plasmid-paired and unpaired discordant sites detected in ARC by WGS and mutant phenotypes.

S2 Table. Mutants with deletions unassociated with plasmid insertion.

S3 Table. Total genes affected in ARC and their description.

S4 Table. Higher-confidence candidate genes and corresponding mutants.

S5 Table. List of PCR primers used in this study.

S1 Fig. Proportion of different types of insertions observed in ARC.

S2 Fig. Genetic linkage test of par^R^ and Ac-phenotypes.

S3 Fig. Two mutant alleles in tocopherol cyclase (Cre01.g013801*, VTE1*) in ARC.

S1 Appendix. Citations from S4 Table.

