## Supplementary material for "Discovery of photosynthesis genes through whole-genome sequencing of acetate-requiring mutants of *Chlamydomonas reinhardtii*": S1 Appendix Table S4 reference list

**Supplemental reference accompanying Table S4**


**Adhikari, N.D., Froehlich, J.E., Strand, D.D., Buck, S.M., Kramer, D.M. and Larkin, R.M.** (2011) GUN4-porphyrin complexes bind the chlH/GUN5 subunit of Mg-chelatase and promote chlorophyll biosynthesis in *Arabidopsis*. *Plant Cell*, **23**, 1449–1467.

**Albee, A.J., Kwan, A.L., Lin, H., Granas, D., Stormo, G.D. and Dutcher, S.K.** (2013) Identification of cilia genes that affect cell-cycle progression using whole-genome transcriptome analysis in chlamydomonas reinhardtti. *G3 Genes, Genomes, Genet.*, **3**, 979–991.

**Auchincloss, A.H., Zerges, W., Perron, K., Girard-Bascou, J. and Rochaix, J.D.** (2002) Characterization of Tbc2, a nucleus-encoded factor specifically required for translation of the chloroplast *psbC* mRNA in *Chlamydomonas reinhardtii*. *J. Cell Biol.*, **157**, 953–962.

**Barneche, F., Winter, V., Crèvecœur, M. and Rochaix, J.D.** (2006) ATAB2 is a novel factor in the signalling pathway of light-controlled synthesis of photosystem proteins. *EMBO J.*, **25**, 5907–5918.

**Bellafiore, S., Barneche, F., Peltler, G. and Rochalx, J.D.** (2005) State transitions and light adaptation require chloroplast thylakoid protein kinase STN7. *Nature*, **433**, 892–895.

**Blume, C., Behrens, C., Eubel, H., Braun, H.P. and Peterhansel, C.** (2013) A possible role for the chloroplast pyruvate dehydrogenase complex in plant glycolate and glyoxylate metabolism. *Phytochemistry*, **95**, 168–176.

**Bohne, A.V., Schwarz, C., Schottkowski, M., Lidschreiber, M., Piotrowski, M., Zerges, W. and Nickelsen, J.** (2013) Reciprocal Regulation of Protein Synthesis and Carbon Metabolism for Thylakoid Membrane Biogenesis. *PLoS Biol.*, **11**.

**Boulouis, A., Raynaud, C., Bujaldon, S., Aznar, A., Wollman, F.A. and Choquet, Y.** (2011) The nucleus-encoded trans-acting factor MCA1 plays a critical role in the regulation of cytochrome *f* synthesis in *Chlamydomonas* chloroplasts. *Plant Cell*, **23**, 333–349.

**Brock, A., Brandt, W. and Dräger, B.** (2008) The functional divergence of short-chain dehydrogenases involved in tropinone reduction. *Plant J.*, **54**, 388–401.

**Brzezowski, P., Schlicke, H., Richter, A., Dent, R.M., Niyogi, K.K. and Grimm, B.** (2014) The GUN4 protein plays a regulatory role in tetrapyrrole biosynthesis and chloroplast-to-nucleus signalling in *Chlamydomonas reinhardtii*. *Plant J.*, **79**, 285–298.

**Cai, W., Ma, J., Chi, W., Zou, M., Guo, J., Lu, C. and Zhang, L.** (2010) Cooperation of LPA3 and LPA2 Is Essential for Photosystem II Assembly in *Arabidopsis*. *Plant Physiol.*, **154**, 109–120.

**Calderon, R.H., García-Cerdán, J.G., Malnoë, A., Cook, R., Russell, J.J., Gaw, C., Dent, R.M., Vitry, C. De and Niyogi, K.K.** (2013) A conserved rubredoxin is necessary for photosystem II accumulation in diverse oxygenic photoautotrophs. *J. Biol. Chem.*, **288**, 26688–26696.

**Cheng, Z., Sattler, S., Maeda, H., Sakuragi, Y., Bryant, D.A. and DellaPenna, D.** (2003) Highly Divergent Methyltransferases Catalyze a Conserved Reaction in Tocopherol and Plastoquinone Synthesis in Cyanobacteria and Photosynthetic Eukaryotes. *Plant Cell*, **15**, 2343–2356.

**Chi, X., Zhang, X., Guan, X., Ding, L., Li, Y., Wang, M., Lin, H. and Qin, S.** (2008) Fatty acid biosynthesis in eukaryotic photosynthetic microalgae: Identification of a microsomal delta 12 desaturase in *Chlamydomonas reinhardtii*. *J. Microbiol.*, **46**, 189–201.

**Cohen, Y., Chitnis, V.P., Nechushtai, R. and Chitnis, P.R.** (1993) Stable assembly of PsaE into cyanobacterial photosynthetic membranes is dependent on the presence of other accessory subunits of photosystem I. *Plant Mol. Biol.*

**Dauvillée, D., Stampacchia, O., Girard-Bascou, J. and Rochaix, J.D.** (2003) Tab2 is a novel conserved RNA binding protein required for translation of the chloroplast *psaB* mRNA. *EMBO J.*, **22**, 6378–6388.

**Davies, J.P., Yildiz, F.H. and Grossman, A.** (1996) Sac1, a putative regulator that is critical for survival of *Chlamydomonas reinhardtii* during sulfur deprivation. *EMBO J.*, **15**, 2150–2159.

**Davies, J.P., Yildiz, F.H. and Grossman, A.R.** (1999) Sac3, an Snf1-like serine/threonine kinase that positively and negatively regulates the responses of *Chlamydomonas* to sulfur limitation. *Plant Cell*, **11**, 1179–1190.

**Depège, N., Bellafiore, S. and Rochaix, J.D.** (2003) Role of chloroplast protein kinase Stt7 in LHCII phosphorylation and state transition in *Chlamydomonas*. *Science*, **299**, 1572–1575.

**Díaz-Troya, S., Pérez-Pérez, M.E., Pérez-Martín, M., Moes, S., Jeno, P., Florencio, F.J. and Crespo, J.L.** (2011) Inhibition of protein synthesis by TOR inactivation revealed a conserved regulatory mechanism of the BiP chaperone in *Chlamydomonas*. *Plant Physiol.*, **157**, 730–41.

**Dobáková, M., Sobotka, R., Tichý, M. and Komenda, J.** (2009) Psb28 protein is involved in the biogenesis of the photosystem II inner antenna CP47 (PsbB) in the cyanobacterium *Synechocystis* sp. PCC 6803. *Plant Physiol.*, **149**, 1076–1086.

**Drop, B., Webber-Birungi, M., Fusetti, F., Kouřil, R., Redding, K.E., Boekema, E.J. and Croce, R.** (2011) Photosystem I of *Chlamydomonas reinhardtii* contains nine light-harvesting complexes (Lhca) located on one side of the core. *J. Biol. Chem.*, **286**, 44878–44887.

**Eberhard, S., Loiselay, C., Drapier, D., Bujaldon, S., Girard-Bascou, J., Kuras, R., Choquet, Y. and Wollman, F.A.** (2011) Dual functions of the nucleus-encoded factor TDA1 in trapping and translation activation of atpA transcripts in *Chlamydomonas reinhardtii* chloroplasts. *Plant J.*, **67**, 1055–1066.

**Eggink, L.L., LoBrutto, R., Brune, D.C., Brusslan, J., Yamasato, A., Tanaka, A. and Hoober, J.K.** (2004) Synthesis of chlorophyll b: Localization of chlorophyllide *a* oxygenase and discovery of a stable radical in the catalytic subunit. *BMC Plant Biol.*, **4**.

**Elvira-Matelot, E., Bardou, F., Ariel, F., Jauvion, V., Bouteiller, N., Masson, I. Le, Cao, J., Crespi, M.D. and Vauchereta, H.** (2015) The nuclear ribonucleoprotein SmD1 interplays with splicing, RNA quality control, and posttranscriptional gene silencing in Arabidopsis. *Plant Cell*, **28**, 426–438.

**Engqvist, M.K.M., Kuhn, A., Wienstroer, J., Weber, K., Jansen, E.E.W., Jakobs, C., Weber, A.P.M. and Maurino, V.G.** (2011) Plant D-2-hydroxyglutarate dehydrogenase participates in the catabolism of lysine especially during senescence. *J. Biol. Chem.*, **286**, 11382–11390.

**Eriksson, M.J. and Clarke, A.K.** (1996) The heat shock protein ClpB mediates the development of thermotolerance in the Cyanobacterium *Synechococcus* sp. strain PCC 7942. *J. Bacteriol.*, **178**, 4839–4846.

**Farah, J., Frank, G., Zuber, H. and Rochaix, J.D.** (1995) Cloning and sequencing of a cDNA clone encoding the photosystem I PsaD subunit from *Chlamydomonas reinhardtii*. *Plant Physiol.*, **107**, 1485–1486.

**Gabilly, S.T., Baker, C.R., Wakao, S., Crisanto, T., Guan, K., Bi, K., Guiet, E., Guadagno, C.R. and Niyogi, K.K.** (2019) Regulation of photoprotection gene expression in *Chlamydomonas* by a putative E3 ubiquitin ligase complex and a homolog of CONSTANS. *Proc. Natl. Acad. Sci. U. S. A.*, **116**, 17556–17562.

**García-Cerdán, J.G., Schmid, E.M., Takeuchi, T., et al.** (2020) Chloroplast Sec14-like 1 (CPSFL1) is essential for normal chloroplast development and affects carotenoid accumulation in *Chlamydomonas*. *Proc. Natl. Acad. Sci. U. S. A.*, **117**, 12452–12463.

**Glanz, S., Jacobs, J., Kock, V., Mishra, A. and Kück, U.** (2012) Raa4 is a *trans*-splicing factor that specifically binds chloroplast *tscA* intron RNA. *Plant J.*, **69**, 421–431.

**Grahl, S., Reiter, B., Gügel, I.L.L., Vamvaka, E., Gandini, C., Jahns, P., Soll, J., Leister, D. and Rühle, T.** (2016) The *Arabidopsis* Protein CGLD11 Is Required for Chloroplast ATP Synthase Accumulation. *Mol. Plant*, **9**, 885–899.

**Gromoff, E.D. von, Alawady, A., Meinecke, L., Grimm, B. and Beck, C.F.** (2008) Heme, a plastid-derived regulator of nuclear gene expression in *Chlamydomonas*. *Plant Cell*, **20**, 552–567.

**Hall, M., Kieselbach, T., Sauer, U.H. and Schröder, W.P.** (2012) Purification, crystallization and preliminary X-ray analysis of PPD6, a PsbP-domain protein from *Arabidopsis thaliana*. *Acta Crystallogr. Sect. F Struct. Biol. Cryst. Commun.*, **68**, 278–280.

**Heinnickel, M., Kim, R.G., Wittkopp, T.M., Yang, W., Walters, K.A., Herbert, S.K. and Grossman, A.R.** (2016) Tetratricopeptide repeat protein protects photosystem I from oxidative disruption during assembly. *Proc. Natl. Acad. Sci. U. S. A.*, **113**, 2774–2779.

**Heinnickel, M.L., Alric, J., Wittkopp, T., Yang, W., Catalanotti, C., Dent, R., Niyogi, K.K., Wollman, F.A. and Grossman, A.R.** (2013) Novel thylakoid membrane GreenCut protein CPLD38 impacts accumulation of the cytochrome $b_6f$ complex and associated regulatory processes. *J Biol Chem*, **288**, 7024–7036.

**Hertle, A.P., García-Cerdán, J.G., Armbruster, U., Shih, R., Lee, J.J., Wong, W. and Niyogi, K.K.** (2020) A Sec14 domain protein is required for photoautotrophic growth and chloroplast vesicle formation in *Arabidopsis thaliana*. *Proc. Natl. Acad. Sci. U. S. A.*, **117**, 9101–9111.

**Houille-Vernes, L., Rappaport, F., Wollman, F.A., Alric, J. and Johnson, X.** (2011) Plastid terminal oxidase 2 (PTOX2) is the major oxidase involved in chlororespiration in *Chlamydomonas*. *Proc. Natl. Acad. Sci. U. S. A.*, **108**, 20820–20825.

**Hutin, C., Nussaume, L., Moise, N., Moya, I., Kloppstech, K. and Havaux, M.** (2003) Early light-induced proteins protect *Arabidopsis* from photooxidative stress. *Proc. Natl. Acad. Sci. U. S. A.*, **100**, 4921–4926.

**Inoue, S., Ejima, K., Iwai, E., Hayashi, H., Appel, J., Tyystjarvi, E., Murata, N. and Nishiyama, Y.** (2011) Protection by α-tocopherol of the repair of photosystem II during photoinhibition in *Synechocystis* sp. PCC 6803. *Biochim Biophys Acta*, **1807**, 236–241.

**Ishihara, S., Takabayashi, A., Ido, K., Endo, T., Ifuku, K. and Sato, F.** (2007) Distinct functions for the two PsbP-like proteins PPL1 and PPL2 in the chloroplast thylakoid lumen of *Arabidopsis*. *Plant Physiol.*, **145**, 668–679.

**Ishikawa, Y., Schröder, W.P. and Funk, C.** (2005) Functional analysis of the PsbP-like protein (sll1418) in *Synechocystis* sp. PCC 6803. *Photosynth. Res.*, **84**, 257–262.

**Jensen, P.E., Gibson, L.C.D., Henningsen, K.W. and Hunter, C.N.** (1996) Expression of the chlI, chlD, and chlH genes from the cyanobacterium *Synechocystis* PCC6803 in *Escherichia coli* and demonstration that the three cognate proteins are required for magnesium-protoporphyrin chelatase activity. *J. Biol. Chem.*, **271**, 16662–16667.

**Johnson, X., Steinbeck, J., Dent, R.M., et al.** (2014) Proton Gradient Regulation 5-Mediated

Cyclic Electron Flow under ATP- or Redox-Limited Conditions: A Study of *deltaATPase pgr5* and *deltarbcL pgr5* Mutants in the Green Alga *Chlamydomonas reinhardtii*. *Plant Physiol.*, **165**, 438–452.

**Kanno, T., Venhuizen, P., Wen, T.N., Lin, W.D., Chiou, P., Kalyna, M., Matzke, A.J.M. and Matzke, M.** (2018) PRP4KA, a putative spliceosomal protein kinase, is important for alternative splicing and development in *Arabidopsis thaliana*. *Genetics*, **210**, 1267–1285.

**Khrebtukova, I. and Spreitzer, R.J.** (1996) Elimination of the *Chlamydomonas* gene family that encodes the small subunit of ribulose-1,5-bisphosphate carboxylase/oxygenase. *Proc Natl Acad Sci U S A*, **93**, 13689–13693.

**Kinoshita, A., Betsuyaku, S., Osakabe, Y., et al.** (2010) RPK2 is an essential receptor-like kinase that transmits the CLV3 signal in *Arabidopsis*. *Development*, **137**, 3911–3920.

**Knoppová, J., Yu, J., Konik, P., Nixon, P.J. and Komenda, J.** (2016) CyanoP is involved in the early steps of Photosystem II assembly in the cyanobacterium *Synechocystis* sp. PCC 6803. *Plant Cell Physiol.*, **57**, 1921–1931.

**Krieger-Liszkay, A. and Feilke, K.** (2016) The dual role of the plastid terminal oxidase PTOX: Between a protective and a pro-oxidant function. *Front. Plant Sci.*, **6**.

**Krieger-Liszkay, A., Shimakawa, G. and Sétif, P.** (2020) Role of the two PsaE isoforms on $O_2$ reduction at photosystem I in *Arabidopsis thaliana*. *Biochim. Biophys. Acta - Bioenerg.*, **1861**.

**Kubo, T., Yagi, T. and Kamiya, R.** (2012) Tubulin polyglutamylation regulates flagellar motility by controlling a specific inner-arm dynein that interacts with the dynein regulatory complex. *Cytoskeleton*, **69**, 1059–1068.

**Kuras, R., Saint-Marcoux, D., Wollman, F.A. and Vitry, C. De** (2007) A specific *c*-type cytochrome maturation system is required for oxygenic photosynthesis. *Proc. Natl. Acad. Sci. U. S. A.*, **104**, 9906–9910.

**Larkin, R.M., Alonso, J.M., Ecker, J.R. and Chory, J.** (2003) GUN4, a regulator of chlorophyll synthesis and intracellular signaling. *Science*, **299**, 902–906.

**Leborgne-Castel, N., Jelitto-Van Dooren, E.P.W.M., Crofts, A.J. and Denecke, J.** (1999) Overexpression of BiP in tobacco alleviates endoplasmic reticulum stress. *Plant Cell*, **11**, 459–469.

**Lefebvre-Legendre, L., Choquet, Y., Kuras, R., Loubéry, S., Douchi, D. and Goldschmidt-Clermont, M.** (2015) A nucleus-encoded chloroplast protein regulated by iron availability governs expression of the photosystem I subunit PsaA in *Chlamydomonas reinhardtii*. *Plant Physiol.*, **167**, 1527–1540.

**Lennartz, K., Plücken, H., Seidler, A., Westhoff, P., Bechtold, N. and Meierhoff, K.** (2001) HCF164 encodes a thioredoxin-like protein involved in the biogenesis of the cytochrome *b6f* complex in *Arabidopsis*. *Plant Cell*, **13**, 2539–2551.

**Lezhneva, L., Kuras, R., Ephritikhine, G. and Vitry, C. De** (2008) A novel pathway of cytochrome *c* biogenesis is involved in the assembly of the cytochrome $b_6f$ complex in *Arabidopsis* chloroplasts. *J. Biol. Chem.*, **283**, 24608–24616.

**Li, H.H., Quinn, J., Culler, D., Girard-Bascou, J. and Merchant, S.** (1996) Molecular genetic analysis of plastocyanin biosynthesis in *Chlamydomonas reinhardtii*. *J. Biol. Chem.*, **271**, 31283–31289.

**Li, X., Patena, W., Fauser, F., et al.** (2019) A genome-wide algal mutant library and functional

screen identifies genes required for eukaryotic photosynthesis. *Nat. Genet.*, **51**, 627–635.
**Liu, J., Yang, H., Lu, Q., Wen, X., Chen, F., Peng, L., Zhang, L. and Lu, C.** (2013) PSBP-DOMAIN PROTEIN1, a Nuclear-Encoded thylakoid lumenal protein, is essential for photosystem I assembly in *Arabidopsis*. *Plant Cell*, **24**, 4992–5006.
**Liu, X.L., Yu, H.D., Guan, Y., Li, J.K. and Guo, F.Q.** (2012) Carbonylation and loss-of-function analyses of SBPase reveal its metabolic interface role in oxidative stress, carbon assimilation, and multiple aspects of growth and development in arabidopsis. *Mol. Plant*, **5**, 1082–1099.
**Mackinder, L.C.M., Chen, C., Leib, R.D., Patena, W., Blum, S.R., Rodman, M., Ramundo, S., Adams, C.M. and Jonikas, M.C.** (2017) A Spatial Interactome Reveals the Protein Organization of the Algal $CO_2$-Concentrating Mechanism. *Cell*, **171**, 133-147.e14.
**Moll, B. and Levine, R.P.** (1970) Characterization of a Photosynthetic Mutant Strain of *Chlamydomonas reinhardi* Deficient in Phosphoribulokinase Activity. *Plant Physiol.*, **46**, 576–580.
**Mozzo, M., Mantelli, M., Passarini, F., Caffarri, S., Croce, R. and Bassi, R.** (2010) Functional analysis of Photosystem I light-harvesting complexes (Lhca) gene products of *Chlamydomonas reinhardtii*. *Biochim. Biophys. Acta - Bioenerg.*, **1797**, 212–221.
**Nath, K., Wessendorf, R.L. and Lu, Y.** (2016) A nitrogen-fixing subunit essential for accumulating 4Fe-4S-containing photosystem I core proteins. *Plant Physiol.*, **172**, 2459–2470.
**Ozawa, S.I., Bald, T., Onishi, T., Xue, H., Matsumura, T., Kubo, R., Takahashi, H., Hippler, M. and Takahashi, Y.** (2018) Configuration of Ten Light-Harvesting Chlorophyll *a/b* Complex I Subunits in *Chlamydomonas reinhardtii* Photosystem I. *Plant Physiol.*, **178**, 583–595.
**Perlaza, K., Toutkoushian, H., Boone, M., Lam, M., Iwai, M., Jonikas, M.C., Walter, P. and Ramundo, S.** (2019) The Mars1 kinase confers photoprotection through signaling in the chloroplast unfolded protein response. *Elife*, **8**.
**Porfirova, S., Bergmüller, E., Tropf, S., Lemke, R. and Dörmann, P.** (2002) Isolation of an *Arabidopsis* mutant lacking vitamin E and identification of a cyclase essential for all tocopherol biosynthesis. *Proc. Natl. Acad. Sci. U. S. A.*, **99**, 12495–12500.
**Pružinská, A., Tanner, G., Anders, I., Roca, M. and Hörtensteiner, S.** (2003) Chlorophyll breakdown: Pheophorbide a oxygenase is a Rieske-type iron-sulfur protein, encoded by the *accelerated cell death 1 gene*. *Proc. Natl. Acad. Sci. U. S. A.*, **100**, 15259–15264.
**Rivier, C., Goldschmidt-Clermont, M. and Rochaix, J.D.** (2001) Identification of an RNA-protein complex involved in chloroplast group II intron trans-splicing in *Chlamydomonas reinhardtii*. *EMBO J.*, **20**, 1765–1773.
**Rojas-González, J.A., Soto-Súarez, M., García-Díaz, Á., et al.** (2015) Disruption of both chloroplastic and cytosolic FBPase genes results in a dwarf phenotype and important starch and metabolite changes in *Arabidopsis thaliana*. *J. Exp. Bot.*, **66**, 2673–2689.
**Roose, J.L., Frankel, L.K. and Bricker, T.M.** (2014) The PsbP domain protein 1 functions in the assembly of lumenal domains in photosystem I. *J. Biol. Chem.*, **289**, 23776–23785.
**Sahrawy, M., Ávila, C., Chueca, A., Cánovas, F.M. and López-Gorgé, J.** (2004) Increased sucrose level and altered nitrogen metabolism in *Arabidopsis thaliana* transgenic plants expressing antisense chloroplastic fructose-1,6-bisphosphatase. *J. Exp. Bot.*, **55**, 2495–2503.
**Schneidere, D., Berry, S., Volkmer, T., Seidler, A. and Rögner, M.** (2004) PetC1 is the major

Rieske iron-sulfur protein in the cytochrome *b₆f* complex of *Synechocystis* sp. PCC 6803. *J. Biol. Chem.*, **279**, 39383–39388.
**Serrano, G., Herrera-Palau, R., Romero, J.M., Serrano, A., Coupland, G. and Valverde, F.** (2009) Chlamydomonas *CONSTANS* and the Evolution of Plant Photoperiodic Signaling. *Curr. Biol.*, **19**, 359–368.
**Shintani, D.K., Cheng, Z. and DellaPenna, D.** (2002) The role of 2-methyl-6-phytylbenzoquinone methyltransferase in determining tocopherol composition in *Synechocystis* sp. PCC6803. *FEBS Lett.*, **511**, 1–5.
**Shtaida, N., Khozin-Goldberg, I., Solovchenko, A., Chekanov, K., Didi-Cohen, S., Leu, S., Cohen, Z. and Boussiba, S.** (2014) Downregulation of a putative plastid PDC E1α subunit impairs photosynthetic activity and triacylglycerol accumulation in nitrogen-starved photoautotrophic *Chlamydomonas reinhardtii*. *J. Exp. Bot.*, **65**, 6563–6576.
**Somanchi, A., Barnes, D. and Mayfield, S.P.** (2005) A nuclear gene of *Chlamydomonas reinhardtii, Tba1*, encodes a putative oxidoreductase required for translation of the chloroplast psbA mRNA. *Plant J.*, **42**, 341–352.
**Soupene, E., Inwood, W. and Kustu, S.** (2004) Lack of the Rhesus protein Rh1 impairs growth of the green alga *Chlamydomonas reinhardtii* at high $CO_2$. *Proc. Natl. Acad. Sci. U. S. A.*, **101**, 7787–7792.
**Takada, S., Wilkerson, C.G., Wakabayashi, K.I., Kamiya, R. and Witman, G.B.** (2002) The outer dynein arm-docking complex: Composition and characterization of a subunit (Oda1) necessary for outer arm assembly. *Mol. Biol. Cell*, **13**, 1015–1029.
**Thornton, L.E., Ohkawa, H., Roose, J.L., Kashino, Y., Keren, N. and Pakrasi, H.B.** (2004) Homologs of plant PsbP and PsbQ proteins are necessary for regulation of photosystem II activity in the cyanobacterium *Synechocystis* 6803 W inside box sign. *Plant Cell*, **16**, 2164–2175.
**Tokutsu, R., Fujimura-Kamada, K., Matsuo, T., Yamasaki, T. and Minagawa, J.** (2019) The CONSTANS flowering complex controls the protective response of photosynthesis in the green alga *Chlamydomonas*. *Nat. Commun.*, **10**.
**Walters, R.G.** (2003) Identification of Mutants of Arabidopsis Defective in Acclimation of Photosynthesis to the Light Environment. *Plant Physiol.*, **131**, 472–481.
**Wang, F., Johnson, X., Cavaiuolo, M., Bohne, A.V., Nickelsen, J. and Vallon, O.** (2015) Two *Chlamydomonas* OPR proteins stabilize chloroplast mRNAs encoding small subunits of photosystem II and cytochrome *b₆f*. *Plant J.*, **82**, 861–873.
**Wittkopp, T.M., Saroussi, S., Yang, W., et al.** (2018) GreenCut protein CPLD49 of *Chlamydomonas reinhardtii* associates with thylakoid membranes and is required for cytochrome *b₆f* complex accumulation. *Plant J.*, **94**, 1023–1037.
**Xing, J., Liu, P., Zhao, L. and Huang, F.** (2017) Deletion of CGLD1 impairs PSII and increases singlet oxygen tolerance of green alga *Chlamydomonas reinhardtii*. *Front. Plant Sci.*, **8**, 2154.
**Xu, Q., Armbrust, T.S., Guikema, J.A. and Chitnis, P.R.** (1994) Organization of Photosystem I polypeptides. A structural interaction between the PsaD and PsaL subunits. *Plant Physiol.*, **106**, 1057–1063.
**Yamano, T., Tsujikawa, T., Hatano, K., Ozawa, S.I., Takahashi, Y. and Fukuzawa, H.** (2010) Light and low-$CO_2$-dependent LCIBLCIC complex localization in the chloroplast supports the

carbon-concentrating mechanism in *Chlamydomonas reinhardtii*. *Plant Cell Physiol.*, **51**, 1453–1468.

**Ying, Z., Mulligan, R.M., Janney, N. and Houtz, R.L.** (1999) Rubisco small and large subunit N-methyltransferases. Bi- and mono- functional methyltransferases that methylate the small and large subunits of Rubisco. *J. Biol. Chem.*, **274**, 36750–36756.

**Yoshihara, C., Inoue, K., Schichnes, D., Ruzin, S., Inwood, W. and Kustu, S.** (2008) An Rh1-GFP fusion protein is in the cytoplasmic membrane of a white mutant strain of *Chlamydomonas reinhardtii*. *Mol. Plant*, **1**, 1007–1020.
